# Single-base 5mC and 5hmC profiling identifies a dynamic 5hmC-to-5mC transition at gene bodies and transcription end sites during neuronal differentiation of mouse embryonic stem cells

**DOI:** 10.64898/2026.11.20.768104

**Authors:** Daniel J. Evanich, Laura N. Blum, Sagnik Sen, Matthew A. Campbell, Louise Williams, Bradley W. Langhorst, V K Chaithanya Ponnaluri, Sriharsa Pradhan

## Abstract

5-methylcytosine (5mC) and 5-hydroxymethylcytosine (5hmC) are genome-wide epigenetic modifications involved in chromatin organization and transcriptional regulation. Although Enzymatic Methyl-seq (EM-seq) can enable the combined profiling of 5mC and 5hmC, Enzymatic 5hmC-seq (E5hmC-seq) can provide selective profiling of 5hmC at single-base resolution. In this context, we systematically validated E5hmC-seq using diverse genomic DNA samples. Next, we integrated E5hmC-seq with EM-seq to distinguish between 5mC and 5hmC landscapes during neuronal differentiation of mouse embryonic stem cells. First, we observed dynamic remodeling of both modifications throughout differentiation, with pronounced changes during the transition from embryoid bodies (EBs) to neural progenitor cells (NPCs), characterized by a global decrease in 5hmC accompanied by an increase in 5mC. Subsequently, differentially hydroxymethylated regions (DhMRs) exhibiting 5hmC depletion frequently showed a concurrent increase in 5mC, whereas regions gaining 5hmC displayed an opposing trend, reflecting the genome-wide relationship. Mapping these modifications to transcriptional profiles revealed an association between cytosine modification dynamics and gene expression, where transcriptional downregulation correlated with 5hmC depletion with concomitant 5mC enrichment. In summary, combined mapping of EM-seq and E5hmC-seq can decipher the dynamic interplay between 5mC and 5hmC at single-base resolution and provide insights into the relationships among cytosine modification dynamics, genomic context and transcriptional regulation during differentiation.

## Introduction

In vertebrate genomes, epigenetic modification is crucial for maintaining gene expression and determining cell fate. Complex epigenetic states can be induced by several converging and amplifying cellular signals and factors, including chromatin associated proteins (CAPs), non-coding RNAs (ncRNA), RNA modifications, DNA-methylation, and post translational modifications (PTM) of histones (1-5). All these processes are dynamic, reversible, and orchestrated throughout cell cycle and development. The most widely studied epigenetic modification mechanism in mammals is DNA methylation (6). It occurs by the covalent attachment of a methyl group to the 5th carbon of cytosine by a family of DNA (cytosine-5) methyltransferases: DNMT1, DNMT3A, DNMT3B using S-adenosyl-L-methionine cofactor (7-10). There are also several accessory factors that guide the methyltransferases to their target sites, namely PCNA and UHRF1 (11,12). It is commonly accepted that when methylation occurs near a promoter sequence, the gene is generally repressed, because proteins that bind to the methylated CG prevents transcription factors from binding to DNA. Conversely, accessible regions in the genome are transcriptionally active and are free of CG methylation (13,14). This hypothesis was revised after the discovery of 5-methylcytosine (5mC) oxidase, ten-eleven translocation (TET) families. TET enzymes with α-ketoglutarate (α-KG), and molecular oxygen would participate in successive oxidation of 5mC to 5hmC (5-hydroxymethylcytosine), 5fC (5-formylcytosine) and 5caC (5-carbxycytosine) (15,16).

5hmC is a relatively rare DNA modification compared to its precursor 5mC and is found across the genome (17). Genome-wide, 5hmC is more abundant at regulatory regions near transcription start sites (TSSs), enhancers, promoters, and exons, consistent with its role in gene expression regulation (18,19). The levels of 5hmC are enriched at active promoter regions, as observed upon activation of neuronal function-related genes in neural progenitors and neurons (20,21). Furthermore, 5hmC weakens the interaction between DNA and nucleosomal H2A-H2B dimers, decreasing thermodynamic stability of the DNA duplex, and facilitating RNA polymerase II to participate in the elongation process (22). Analysis of genome-wide 5hmC distribution profiles shows a positive correlation between 5hmC and R-loops in mouse embryonic stem (mES), and in human embryonic kidney 293 (HEK293) cells suggesting its role in transcription (23). Oxidation of 5mC results in active demethylation of the genome initiated through the progressive oxidation of 5mC to 5hmC, 5fC, and 5caC (24). This process sets the stage for DNA demethylation by one of the two potential pathways. First, 5hmC is not well recognized by the DNMT1 maintenance methylation machinery allowing for its passive loss over several rounds of DNA replication correlating with cell division (25). Secondly, 5fC and 5caC can be targeted for base excision by thymine DNA glycosylase (TDG) (26-28). This 5mC erasure is particularly important in pluripotent embryonic stem cells (ESCs) to maintain self-renewal and retain the potential to generate different cell types with diverse 5mC patterns following epigenetic reprogramming. In fact, elevated levels of 5hmC observed in embryonic stem cells and neuronal cells appeared to correlate with pluripotency and neuronal development (5). In our previous study, we have demonstrated a gradual increase in the total 5mC in conjunction with a rapid increase followed by a slow and gradual decrease of 5hmC levels during embryoid body differentiation in mouse E14 stem cells (29). Furthermore, genome wide studies using hmeDIP-seq have demonstrated 5hmC modifications enriched in gene bodies and enhancers as a signature mark for transcriptional activation (19). These studies relied on 5hmC enrichment via immunoprecipitation or differential restriction enzyme digestion to quantify 5hmC, which lacks true single base resolution.

With the development of technologies to map DNA modifications genome wide, it is now possible to map 5hmC at single base resolution for biological studies (30). One notable method is TET-assisted bisulfite sequencing method (TAB-seq) that is based on 5hmCs in genomic DNA being protected by glucosylation to make 5gmC, and then 5mCs are successively converted to 5caCs by enzymatic oxidation of TET. After bisulfite treatment, both 5caC (generated from TET mediated oxidation of 5mC) and unmodified C display as T while 5gmC (generated from glucosylation of original 5hmC) display as C (31). However, it is known that bisulfite treatment can lead to significant degradation of input DNA by as much as 99% (32). It is possible to replace bisulfite with APOBEC and enzymatically convert all C and 5mC to U and T that is read as T following amplification, as demonstrated by E5hmC-seq and APOBEC-Coupled Epigenetic Sequencing (33). Despite all available mapping technologies, a systematic study of the dynamics between 5mC and 5hmC at single base resolution and their interplay for transcriptional gene regulation is lacking.

In this report, we have extensively validated and presented the results for an enzymatic method that specifically detects 5hmC at single base resolution genome wide using Enzymatic 5hmC-seq (E5hmC-seq, NEB). We rigorously profiled 5hmC distribution in the genome of various mouse tissues to demonstrate tissue specific 5mC and 5hmC distribution and the associated pathways. In addition, we have used mouse E14 embryonic stem cells to neuronal cell differentiation as a model system to investigate the interplay between 5mC and 5hmC dynamics. A subtractive analysis of genome wide methylation marks using EM-seq (5mC and 5hmC) (34) and E5hmC-seq (5hmC) allowed us to differentiate both epigenetic marks. Finally, we performed detailed analysis of differentially methylated regions for 5mC and 5hmC and their correlation with transcription was undertaken for selected genes.

## Results

### Enzymatic 5hmC-seq (E5hmC-seq) of gDNA

The E5hmC-seq method relies on two enzymatic reactions on genomic DNA (gDNA) to detect 5hmC at single base resolution. The first step is glucosylation of 5hmC using T4 Phage β-glucosyltransferase (T4-BGT), which transfers the glucose moiety to 5hmC from uridine diphosphoglucose (UDP-Glucose) to generate β-glucosyl-5-hydroxymethylcytosine (β-g5hmC). In the second step, cytosine and 5mC are deaminated to uracil and thymine, respectively by apolipoprotein B mRNA editing enzyme catalytic subunit 3A (APOBEC3A) leaving β-g5hmC formed in the prior step. After PCR amplification, the deaminated C and 5mC residues are read as T and the β-g5hmC as C (Figure 1A).

**Figure 1.**
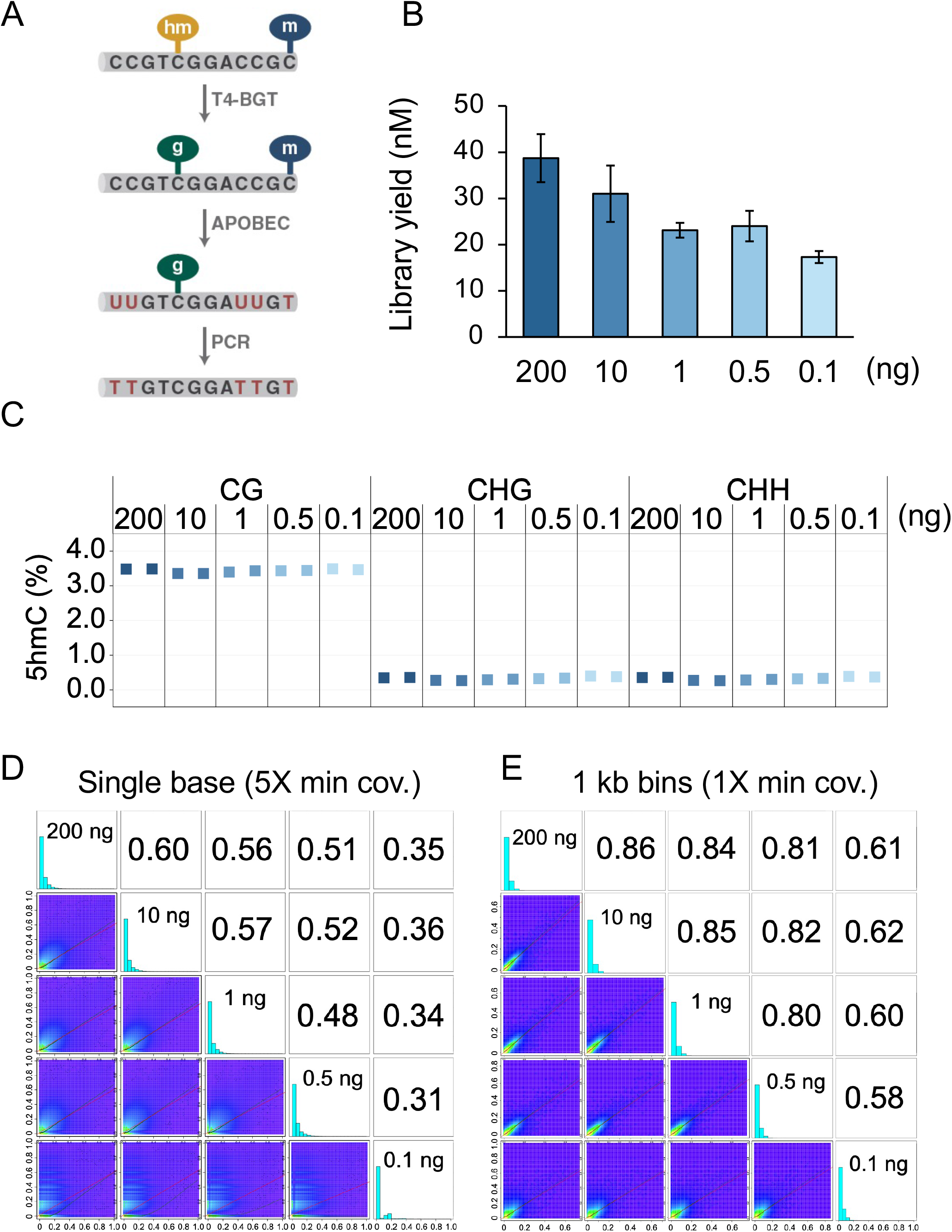
E5hmC-seq and library metrics across various amounts of gDNA. (A) A schematic diagram for specific 5hmC detection is shown. 5hmC is first glucosylated using T4-BGT followed by 5mC and unmodified cytosine deamination by APOBEC to thymine and uracil, respectively. During NextGen sequencing, 5hmCs are represented as cytosine, while cytosine and 5mCs are represented as thymine. (B) Library yields for E5hmC-seq libraries prepared from 200 – 0.1 ng of mouse E14 genomic DNA is shown. Yields shown are the average of two technical replicates per library. (C) 5hmC levels in undifferentiated E14 mouse embryonic stem cell gDNA in the CG, CHG, and CHH contexts is shown. Approximately 3.5% of 5hmC in the CG context is detected across all input amounts. (D) Pearson correlations for E5hmC-seq libraries generated using methylKit. Data shown for each input is the combination of two technical replicate libraries. Correlations were generated at single base level for 200 ng – 0.1ng at 5X minimum coverage or (E) after averaging 5hmC levels in 1 kb sliding windows. All CGs included in the windowing analysis are at 1X minimum coverage, and all windows contain at least 5 CG sites.

For detailed evaluation of E5hmC-seq performance, we used different amounts of mouse ES cell E14 gDNA (200 ng, 10 ng, 1 ng, 0.5 ng and 0.1 ng) along with spike-in control DNA (fully hydroxymethylated T4, and fully unmethylated lambda DNAs) to monitor enzymatic conversion. Duplicate libraries were made and sequenced using Illumina Novaseq 6000. E5hmC-seq libraries had high yields across the DNA amounts (Figure 1B). From the aligned reads, we calculated the conversion efficiency to monitor 5hmC levels. We observed ≥ 98.9% of sequenced cytosines in fully hydroxymethylated T4 control DNA across CG, CHG and CHH sequences as 5hmC (Figure S1A). The unmethylated lambda control displayed ≥ 99.5% of cytosine deamination (read as T in sequencing; Figure S1B). Taken together, these results confirm that both glucosylation and deamination reactions are highly efficient and stable regardless of the DNA amount tested in our study. Further, in the E14 genome, aggregate 5hmC level in the CGs was ∼3.5% compared to low background levels at the CHG and CHH sequences across all tested amounts (Figure 1C). Additionally, E5hmC-seq libraries showed a consistent insert size distribution that peaks around 250 bp (Figure. S1C). The E5hmC-seq libraries detected at least 40 million CG sites at 1X depth for the 200 ng to 0.5 ng gDNA and ∼38.7 million CG sites at 1X depth for the 0.1 ng gDNA compared to an estimated ∼44 million CGs present in the grcm39 mouse genome (Figure S1D). Many of those CG sites covered demonstrated high depth of coverage, including 30 million CGs with minimum 25X coverage for 200 and 10 ng samples, and 13.5 million CGs in the 1 ng samples, respectively. Analysis of the GC bias profile showed that E5hmC-seq libraries had even coverage across the genome (Figure S1E). We observed low (< 37%) duplication rates for 200, 10, and 1 ng DNA and higher duplication rates ∼53% for 0.5 ng DNA and ∼83% for 0.1 ng DNA due to higher numbers of PCR cycles. Similarly, the percent usable (properly-paired, MapQ 10+, primary, nonduplicate) reads for methylation calling remained ≥ 55% for 200, 10, and 1 ng DNA; ∼40% for 0.5 ng DNA and over 10% for the lowest amount i.e. 0.1 ng of DNA (Figure S1F). We merged both technical replicates to gain higher genome coverage (Figure S1G).

E5hmC-seq libraries displayed similar number of genomic features at ≥ 5X coverage across different DNA amounts for diverse genomic features till 0.5 ng of gDNA (Figure S1H). Pearson correlations for E5hmC-seq libraries (200, 10, 1 and 0.5 ng) at single base level with 5X minimum coverage showed a correlation coefficient of ∼0.3 or higher. Unlike the bimodal distribution of 5mC at 0 and 100%, the majority of the sites with 5hmC tend to be hydroxymethylated at a low level with very few exceptions at 100% (Figure 1D). By averaging the 5hmC level across 1kb bins in the genome, we observed a higher Pearson correlation coefficient of ∼0.85 between 200, 10, 1, and 0.5 ng DNA samples along with an improvement in the correlation for 0.1 ng DNA (Figure 1E). Finally, the 5hmC levels detected in T4 spike in control across all three sequence contexts were highly correlated with a correlation coefficient of (≥ 0.9) at single base level demonstrating consistent detection of the modified 5hmC base using E5hmC-seq (Figure S1I-K).

### Comparison between hMeDIP-seq and E5hmC-seq

Since hMeDIP has been used extensively to identify regions in the genome enriched for 5hmC modification, we compared hMeDIP-seq with E5hmC-seq for the human brain gDNA. Regions in hMeDIP which were significantly enriched were identified by calling peaks against the input control. We then checked these enriched regions for CG sites covered in our deep-sequenced 5hmC-seq dataset. The false positive rate was low, with only around 2% of peaks showing no detectable 5hmC (i.e., at least one site with 5hmC above 0%) (Figure S1L). We binned CG sites according to their 5hmC level and found that of the 53M CGs which were covered with at least 5X in E5hmC-seq, around 13M had 0% 5hmC. Of the sites containing 5hmC, we saw that the proportion of CGs enriched by hMeDIP increased with 5hmC level. Of the 7.6M CGs with the highest 5hmC level (over 40%), 38% were significantly enriched (Figure S1M). This means a majority (62%) of the highest-5hmC CGs were not significantly enriched by hMeDIP, representing false negatives. Among CG sites with lower 5hmC levels, e.g., less than 10% of 5hmC, only around 15% were significantly enriched. The observation that hMeDIP enrichment is positively correlated to 5hmC level confirms that the method is robust. However, the majority of 5hmC-containing CGs detected by 5hmC-seq are not significantly enriched by hMeDIP, which suggests that E5hmC-seq offers a much higher degree of sensitivity for assessing even the lowest 5hmC levels.

### Enzymatic 5hmC-seq accurately predicts global levels of 5hmC in mouse tissue

To measure 5hmC modifications globally, we analyzed mouse gDNA extracted from 14 different tissues of origin: brain, colon, heart, intestine, kidney, liver, lung, pancreas, skeletal muscle, skin, spleen, stomach, placenta, and uterus. We generated E5hmC-seq and EM-seq libraries from two biological replicates gDNA samples per tissue and sequenced all libraries to an average of ∼20 million total reads per sample. Analysis of spike-in controls in these libraries confirmed the efficiency of the enzymatic conversion reactions across both E5hmC-seq and EM-seq library types (Figures S1N-S). The mapping rate was consistent across all samples (∼99.5% on average), with low duplication levels (∼8% on average), for both E5hmC-seq and EM-seq libraries (Supplementary Table S1). E5hmC-seq revealed high diversity in 5hmC levels at CG sites across different tissues that ranged from 12.4% in brain down to 1.4% in spleen (Figure 2A). There were low amounts of 5hmC in the non-CG context for all tissue gDNAs (Figure S2A). CG methylation (both 5mC and 5hmC) was consistent across all tissues (between 70-78%) except for placenta where the level was 55.4% (Figure 2B). As expected, methylation levels were low in the non-CG sequence context, except for brain with ∼1% methylation (Figure S2B). Additionally, to validate the sequencing-based levels of 5mC and 5hmC in the genome, we performed LC-MS/MS based global quantification. The 5mC levels remain similar across tissues at 4-5% 5mC of total C, except in placenta where the level was 3.4% (Figure. 2C). In contrast, 5hmC showed greater tissue-specific variation ranging from 1.2% 5hmC of total 5mC in spleen to 11.9% in brain (Figure. 2C). Taken together, here we demonstrate that both EM-seq and E5hmC-seq can profile cytosine methylation and hydroxymethylation accurately and reproducibly genome-wide at single base resolution from genomic DNA. The diverse range of 5hmC levels across different tissues indicates a role for 5hmC in cell fate determination during early embryonic differentiation into the three embryonic germ layers that gives rise to tissue types.

**Figure 2.**
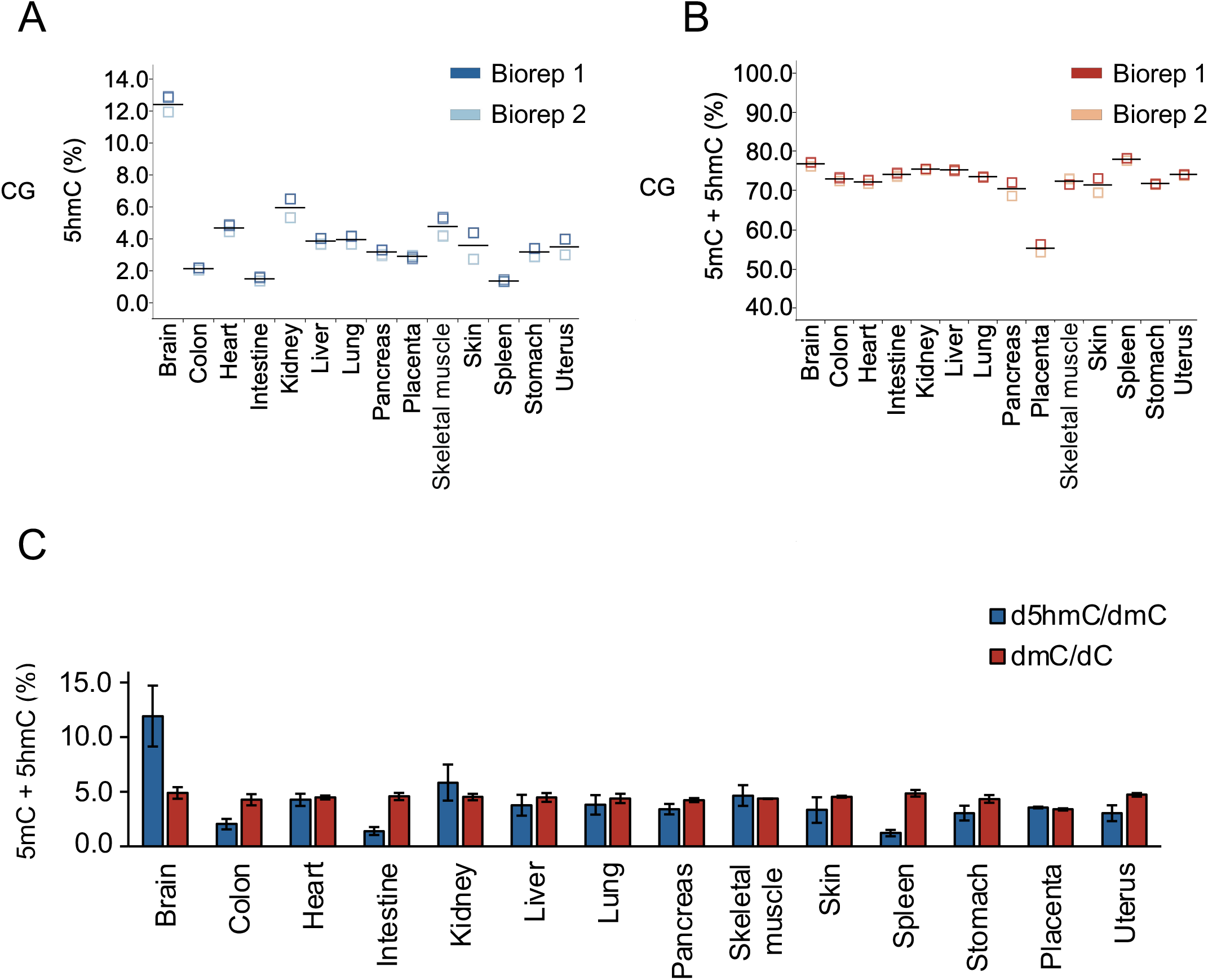
Analysis of 5mC and 5hmC in diverse mouse tissues gDNA. (A) Percent CG 5hmC detected in 14 different mouse tissues using E5hmC-seq. (B) Percent CG methylation (5mC + 5hmC) in the same mouse tissues gDNA are shown using EM-seq. (C) Average 5mC and 5hmC content based on LC-MS/MS quantification. Error bars in (C) are ± standard deviation.

### Inverse correlation between global 5mC and 5hmC during embryoid body formation but not in neuronal differentiation

Previously we have observed a gradual increase in the total 5mC in conjunction with a rapid increase followed by a slow and gradual decrease in 5hmC levels in the genome during ES to embryoid body differentiation (29). Since cytosine hydroxymethylation is abundant in neuronal cells genomic DNA, we hypothesized that ES to neuronal cell differentiation would also involve dynamic interplay of DNA cytosine modification (5mC and 5hmC). We performed a differentiation time course experiment where the cells were transitioned from their pluripotent state to neuronal lineage in a two-step process. The first step involved the formation of embryoid bodies by culturing the cells on non-tissue culture plates in media without LIF for four days. On day 4 retinoic acid was added to initiate neuronal progenitor cell formation. Finally, the cells were transitioned into neuronal lineage by introducing N2 supplement on day 8 followed by N2/B27 supplement on day 10. Cells were harvested everyday as depicted in the experimental schematic followed by extraction of genomic DNA and total RNA (Figure 3A). The genomic DNA from various stages of neuronal differentiation were used for (1) global analysis of 5mC and 5hmC levels at each time point. (2) single base resolution of 5mC and 5hmC genome wide. For global 5mC and 5hmC analysis, LC-MS/MS analysis was performed on the digested nucleosides of the genomic DNA. We observed an increase in the 5mC level from day 0 (6.13%) to day 4 (7.44%) and a subsequent gradual decrease to 4.61% on day 13 (Figure 3B). Interestingly, we also observe a consistent decrease of 5hmC level from day 0 (2.76%) to day 6 (1.75%), followed by a steep drop to 0.76% on day 7 and maintaining a steady level till day 13 (Figure 3C). These observations demonstrate an inverse correlation between 5mC and 5hmC in the genome during neuronal differentiation of ES cells.

**Figure 3.**
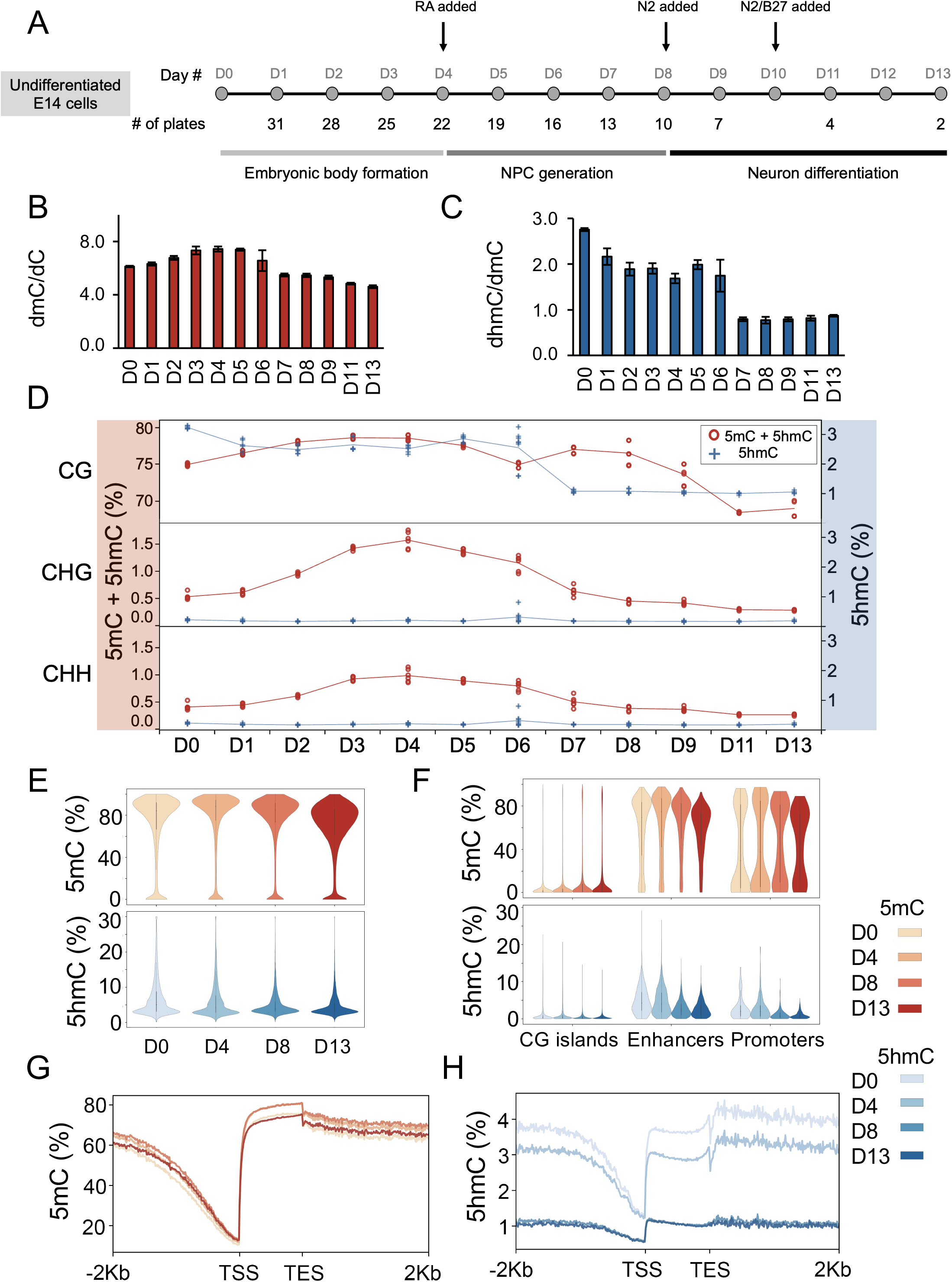
Dynamic methylation and hydroxymethylation during mouse E14 cell gDNA during differentiation. (A) Schematic of the differentiation time course of E14 for neuron differentiation. Undifferentiated cells were split into 31 plates at the beginning of the time course. Each day, three (D1 – D9) or two (D11 and D13) plates were harvested for both RNA and DNA extraction. RA – retinoic acid. Average 5mC (B) and 5hmC (C) content based on LC-MS/MS quantification across the differentiation time course. (D) Mean methylation levels across the time course are represented for EM-seq and E5hmC-seq libraries in CG, CHG, and CHH contexts. Three biological replicates each with two technical replicates are shown. (E-H) For deep-sequenced time points D0, D4, D8, and D13, the 5mC level was calculated by a subtraction between E5hmC-seq and EM-seq libraries. 5mC and 5hmC distributions are represented by kernel density estimates for individual CG sites (E) and for mean methylation levels of CG islands, enhancers, and promoters (F). Metagene plots for 5mC and 5hmC levels in 2kb regions around transcript start and end sites. Error bars in (B) and (C) are ± standard deviation.

To unravel the genomic distribution of the 5mC and 5hmC across the differentiation time-course we prepared EM-seq and E5hmC-seq libraries using biological triplicates and technical replicates. We deep-sequenced libraries for four time points, resulting in an average of 1.6B mapped reads and 67X effective coverage per biological replicate once technical replicates were combined (Supplementary Table S2). We evaluated the biological and technical replicates for consistency in basic metrics like conversion efficiencies of appropriate spike-in controls (Figures S3A-F), GC bias (Figure S3G, H) and insert size distribution (Figure S3I, J). Indeed, we observed a deamination rate of ≥ 99.2% and protection rate of ≥ 99%, along with uniform GC bias and insert size distribution centered around 200-250 bp using both EM-seq and E5hmC-seq libraries. Global aggregate level of 5mC and 5hmC determined using EM-seq showed increasing trends of methylated cytosine in CG from day 0 (75.05%) to day 4 (78.6%) followed by a gradual reduction in modification levels till day 13 (69.03%) (Figure 3D). Interestingly, we also observed an increase in the non-CG methylation from day 0 to day 4 and subsequent reduction to baseline by day 13. However, the 5hmC levels determined by E5hmC-seq showed a steady decline in its levels from day 0 (3.2%) to day 6 (2.4%), followed by a significant drop and stabilization at ∼1% in the CG (Figure 3D). There were no significant differences in the non-CG 5hmC levels unlike previous observation with EM-seq for the same time points. Taken together, we can conclude that the global level of 5mC and 5hmC using LC-MS/MS and our single nucleotide resolution sequencing technology agree with each other.

### Dynamics of 5-methylcytosine and 5-hydroxymethylcytosine at active genomic regions during differentiation

Since the global 5mC and 5hmC levels changed during differentiation and was prominent after addition of retinoic acid (D4) and N2 (D8), we focused on epigenetic transitions and gene expression changes during NPC generation and neuron differentiation. We performed deep sequencing of D0 (stem cell as control), and compared with D4 (embryoid body), D8 (NPC generation), and D13 as the end point of neuron differentiation. Day 4 and 8 cells were perturbed with the addition of retinoic acid, followed by N2 supplements to facilitate transition into NPC, and neuronal differentiation stage, respectively. We evaluated the consistency of CG modification levels across all replicates before combining replicates (Fig. S3K, L).

Using this replicate combined dataset for EM-seq and E5hmC-seq, we performed subtractive analysis to generate 5mC level from the composite (5mC + 5hmC) EM-seq data for D0, 4, 8, and 13. Furthermore, we plotted the distribution of 5mC and 5hmC levels for D0-13 as violin plots. Indeed, the global levels of 5mC and 5hmC showed a gradual decrease genome wide upon differentiation (Figure 3E). Subsequently, we also performed 5mC and 5hmC distribution in different genomic features like CG islands, enhancers, and promoters (Figure 3F). We observe feature type dependent differences in distribution of both 5mC and 5hmC levels with CG islands as expected showing low levels of 5mC and 5hmC modification. Also, enhancers showed a higher level of 5hmC accumulation compared to all other feature types (Figure 3F, Fig S3M). Interestingly, we also observed a high level of 5mC accumulation across the enhancers and bimodal distribution across promoters (Figure 3F).

To understand the distribution of 5mC and 5hmC across TSS and TES we generated a metagene plot with a 2 kb window. We observe an increase in the TSS 5mC level from day 0 to 13. In addition, the gene body 5mC level increased till day 8 followed by a decrease on day 13 (Figure 3G). Interestingly, we observe significantly different 5hmC levels at TSS on day 0 and 4 compared to day 8 and 13. Also, gene body 5hmC level decreased from day 0 to 4, followed by another drop on day 8. This level of 5hmC remained unchanged on day 13 (Figure 3H).

### Developmentally regulated differentially methylated and hydroxymethylated regions during differentiation

Next we interrogated differentially modified regions for 5mC (DMR) and 5hmC (DhMR) across different time points of differentiation to study their interplay and degree of 5mC to 5hmC conversion. We chose day 0-4 (D0-D4) for embryoid body formation, day 0-8 (D0-D8) for NPC, and day 0-13 (D0-D13) for neuronal cell differentiation studies. Using the high coverage 5mC and 5hmC datasets for day 0, 4, 8 and 13 post differentiation, we generated DMR and DhMR calls using day 0 as the control. We identified an increased number of hypo-DhMRs during the differentiation process, D0-D4 (25,533), D0-D8 (225,658), and D0-D13 (223,916) (Figure 4A). As DhMRs accumulated throughout differentiation, most were shared from one comparison to the next (Figure S4A). Over 90% of D0-D4 DhMRs were still detected for D0-D8, and over 80% of DhMRs were shared between D0-D8 and D0-D13. This observation agrees with the global 5hmC trends observed using LC-MS/MS and E5hmC-seq with a sharp drop of 5hmC during day 8 and 13 compared to day 0 (Figure 3C, D). Interestingly, we identified smaller number of hyper-DhMRs during the same differentiation stages, D0-D4 (6,363), D0-D8 (5,541) and D0-D13 (2,613) (Figure 4A). Irrespective of hypo-/hyper methylated DhMR, the magnitude of the median difference in 5hmC level for these DhMRs remained between ∼7-10%.

**Figure 4.**
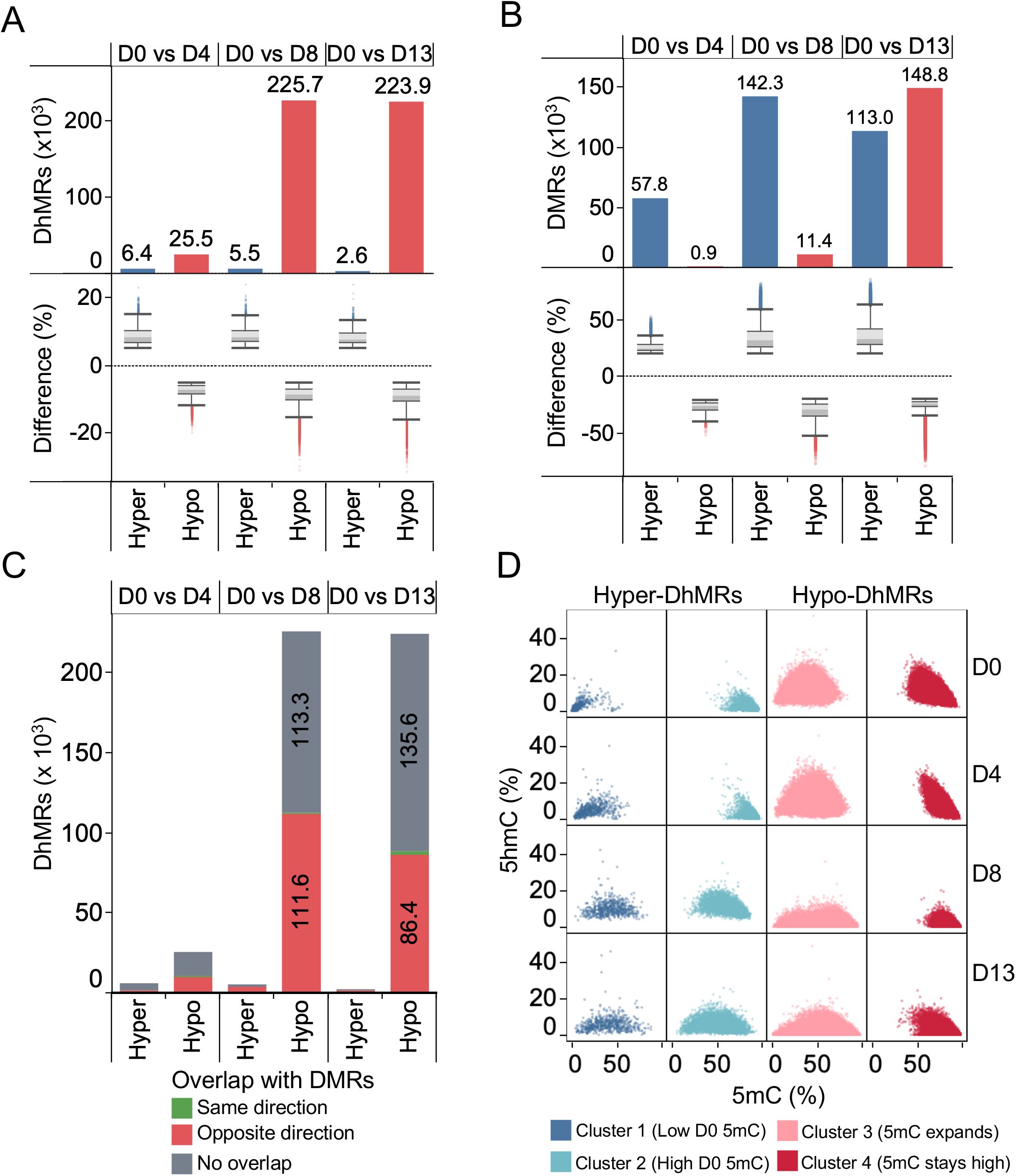
Analysis of differential methylation and hydroxymethyaltion at specific cell linage steps. (A) Number of hyper- and hypo-differentially hydroxymethylated regions (DhMRs) detected for D4, D8, and D13 when compared to D0 with at least 5% change. The distribution of % difference is shown, where boxplot whiskers extend to 1.5xIQR. (B) Number of hyper and hypo differentially methylated regions (DMRs) detected for D4, D8, and D13 when compared to D0 with at least 20% change. The distribution of % difference is shown. (C) The Hyper- and hypo-DHMRs in A classified by overlap with DMRs (minimum 20% overlap), and whether the change in methylation was the same or opposite direction. (D) Mean 5hmC and 5mC methylation levels of hyper- and hypo-DhMRs from the D0 vs D8 comparison across days. DhMRs were clustered based on their 5mC and 5hmC levels across the four time points using k-means clustering, which resulted in clusters 1 and 2 (hyper-DhMRs) and clusters 3 and 4 (hypo-DhMRs)

Next, we performed DMR analysis for 5mC to decipher its link with 5hmC. Using day 0 as the control, we identified hyper-DMRs for D0-D4 (57,848), D0-D8 (142,300), and D0-D13 (148,819) with a median 5mC difference of ∼25% (Figure 4B). Further, D0-D8 and D0-D13 hyper-DMRs showed high degree of overlap (Fig S4B). Interestingly, there was a significant increase in the hypo-DMRs for D0-D13 (148,819) comparison as opposed to D0-D8 (11,380) and D0-D4 (908) (Figure 4B). These results suggest that on D13 there is an equilibrium between hypo- and hypermethylation of 5mC DMRs (149K vs. 110K) in the genome. In addition, there appears to be a correlation between an increased number of hypomethylated DhMRs and hypermethylated DMRs amongst D8 and D13 genome (Figure 4A, B).

Since 5hmC is an oxidative derivative of 5mC and the process is dynamic during development and differentiation, we hypothesized that there will be shared DhMRs and DMRs. We categorized hypo/hyper-DhMRs by analyzing overlapping patterns with hypo/hyper-DMRs (Figure 4C). The distribution of overlapping regions was bimodal: most DhMRs either overlapped a DMR completely or did not overlap (Figure S4A, B). We excluded the least confident overlaps by setting a 20% minimum overlap threshold for a region to be counted as a shared DMR. Interestingly, for all day-to-day comparisons, we noticed ∼50% of DhMRs overlapped a DMR. Furthermore, most of this overlap was hypo-DhMRs with hyper-DMRs, and conversely a small proportion exhibited hyper-DhMRs associated with hypo-DMRs. This suggests these regions are transitioning from one modification state to other (5mC -> 5hmC and 5hmC -> 5mC) between D0-D8 and D0-D13 (Figure 4C).

### Direct transition of 5-hydroxymethylcytosine and 5-methylcytosine during differentiation

To understand the course of DMR/DhMR transition or vice versa over the duration of differentiation time, we performed a trajectory analysis of D0-D8 DhMRs. DhMRs were clustered based on their average 5hmC and 5mC level across the 0, 4, 8 and 13 days (about 2 weeks) of differentiation (Figure 4D). The hyper-DhMRs were separated into two distinct clusters: regions that start with a lower 5mC level that increases over time, along with increasing 5hmC (Cluster 1), and regions that start with a high 5mC level which decreases over the differentiation as 5hmC increases (Cluster 2). For the regions in Cluster 2, a consistent increase in average 5hmC levels along with a decrease in average 5mC level suggests methylation of low 5mC loci followed by TET mediated oxidation to 5hmC (Figure 4D). The hypo-DhMRs can also be described by two clusters. In the first, the average 5mC level on D0 ranges from low to moderately high, and over the time course expands to the edges of the dynamic range as 5hmC decreases (Cluster 3). In the second, 5mC starts out moderately high and shifts higher as 5hmC decreases (Cluster 4). During differentiation, we observe concurrent decreases in 5hmC and increases in 5mC for the same loci, suggesting these sites are transitioning from 5hmC to 5mC perhaps through active demethylation followed by remethylation. Finally, we annotated these DhMR loci by overlap with different genomic features and observed similar trends across different genomic features, (enhancers, exons, lnRNA, mRNA exon, mRNA exon1) suggesting the epigenetic switch is indeed happening across the genic regions (Figure S4C, D).

### Dynamic 5-hydroxymethylcytosine and 5-methylcytosine participate in gene expression

We interrogated the role of DMR and DhMR across different time points of differentiation to study their involvement in transcriptional gene regulation. We performed RNA-seq from D0 to D13 to follow transition from ES to neuronal cell development. The replicates clustered tightly in the PCA plot confirming reproducibility of the RNA-seq data for each sample where PC1 represents variation across the time course (40%) and PC2 represents variation between replicates (17%). Interestingly, samples from D0 to D4 clustered together with D5 and D6 representing transition to tightly clustered time points from D7 to D9, followed by another grouping of D11 and D13 time points. The three distinct clusters represent embryonic body formation, NPC generation, and neuronal differentiation phases of the time course (Figure 5A, Figure 3A). Next, we visualized the trends in average TPM values across the time course as a heatmap with hierarchal clustering. Indeed, few of the transcripts exhibited extreme fluctuation in their average TPM values temporally (Figure S5). To gain a better understanding of these temporal fluctuations in TPM values, we performed differential transcript expression analysis with D0 as control for D4, D8, and D13 time points. We identified high number of significantly differentially expressed transcripts (DETs) for D0-D4: 14596, D0-D8: 19,655, and D0-D13: 25,552 comparisons with a strong p value ≤ 0.05 (Figure 5B, Supplementary Table S3). For D0-D4, 53.9% of DETs were down regulated, whereas 51% and 59% of DETs were up-regulated for NPC and neuronal differentiation stages respectively. Further analysis of up-regulated DETs identified GO: BP terms associated with NPC generation and neuronal differentiation in the D0-D8 and D0-D13 comparisons, respectively (Figure 5C, Supplementary Table S4). GO:BP enrichment results for downregulated DETs are provided in Supplementary Table S5. We also performed a similar analysis using non-differentially expressed transcripts for all three comparisons and observed random enrichment of GO: BP terms with weak p-values (Supplementary Table S6). Next, we analyzed the top 10 GO: BP terms identified using the up-regulated DETs for D0-D13 comparison and plotted the -log10(p-value) across all three comparisons. Our analysis showed a clear increase in the strength of enrichment over the differentiation time course (Figure 5C).

**Figure 5.**
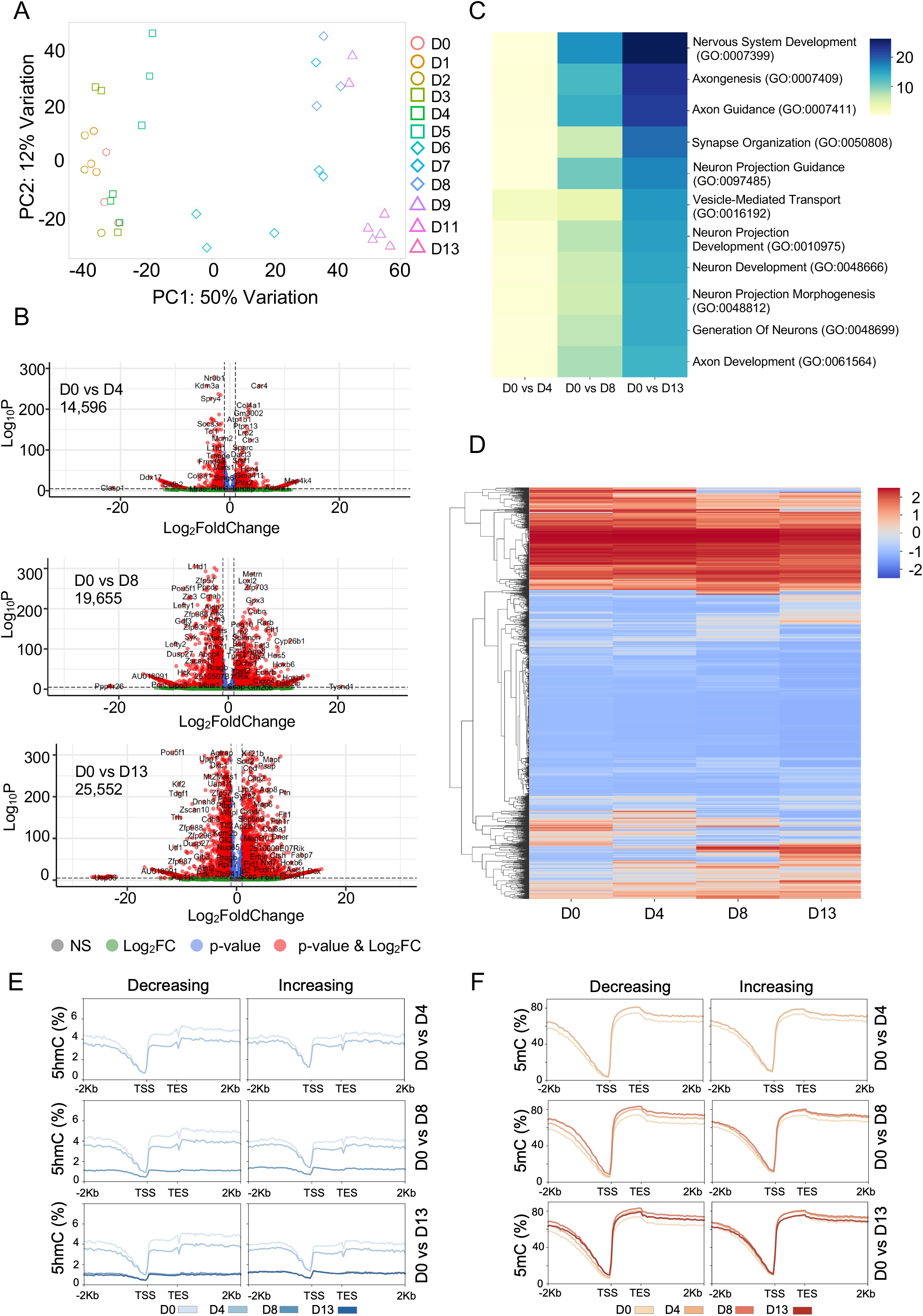
Identification of differential transcript and DNA methylation correlation during E14 differentiation. (A) PCA plot exhibiting the differences between Days (D0-D9, D11, and D13), (B) Volcano plots on DETs to demonstrate the distribution between upregulated DETs and down-regulated DETs for D0vsD4 (total = 14,596 DETs); D0vsD8 (total= 19,655 DETs), and D0vsD13 (total= 25,552 DETs) respectively. (C) List of top GO: BPs (based on -log_10_p-value) associated with D0vsD13 up-regulated DETs and their association with D0vsD4 and D0vsD8 DETs. (D) Distribution of TPM scores for common DETs at D0, D4, D8 and D13 time points. Metagene plot of 5hmC levels (E) and 5mC levels (F) for significantly downregulated and upregulated transcripts for D4, D8, and D13 relative to D0.

Hierarchical clustering analysis across D0, D4, D8, and D13 showed the distribution of TPM for DETs (Figure 5D). DETs were segregated into four classes based on their TPM trajectory: 1. high to high TPM (mostly for down regulated transcripts, TPM dropped significantly compared to D0 however TPM values are high), 2. low to low TPM (mostly for up regulated transcripts, TPM increased significantly compared to D0, however exhibiting relatively low TPM value), 3. high-low TPM (extreme decrease in TPM values) and 4. low-high TPM (extreme increase in TPM values). The size of the hierarchical clusters also exhibited the conservation of all these categories (Figure 5D).

Next, an average of 5hmC and 5mC levels for up and down regulated DETs, we plotted using a metagene plot. We observed significant levels of 5hmC decrease at gene body between D0 and D4 for both up/down regulating transcripts, whereas TSS was not affected (Figure 5E). However, 5hmC enrichment drastically reduced at the TSS in NPC cells (D8) (Figure 5E). In a similar analysis of 5mC levels, we observed a time-dependent increase in the 5mC level at TSS and upstream of TSS from D0 to D8 for down regulated transcripts and subsequent stabilization at D13. We also observed a similar pattern of increasing the 5mC level in the gene body from D0 to D8 followed by a decrease in D13. Interestingly, in the case of up regulated transcripts, we didn’t observe any significant changes in the level of 5mC levels across the metagene plot suggesting these transcripts are regulated in CG methylation independent manner. (Figure 5F).

To understand the relationship between the observed epigenetic changes and transcriptional output, we analyzed enrichment of differential CGs in genes for differentially expressed vs. non-differentially expressed transcripts with respect to 5hmC and 5mC modification during the neuronal cell differentiation process. We used D0 as the control for epigenomic changes and gene expression. Each gene was segmented into the following genomic features: TSS, first exon, internal exons, last exon, first intron, internal introns, last intron, and TES. Each feature type was analyzed separately for either increase or decrease in DNA methylation with transcriptional output. Differential CGs within DMRs and DhMRs intersecting with each feature were counted and normalized to the number of CGs in the feature, followed by a T-test between these values for differentially and non-differentially expressed transcripts (Figure 6, Figure S6, Supplementary Table S7). Our results support the canonical role of 5mC in regulating gene expression, with hyperDMRs in all examined feature types associated with decreased gene expression at D4, D8, and D13 compared to D0 (Figure 6). This observation was further supported by a strong positive correlation between TSS and gene body hypoDMRs in transcriptional upregulation until day 8.

**Figure 6.**
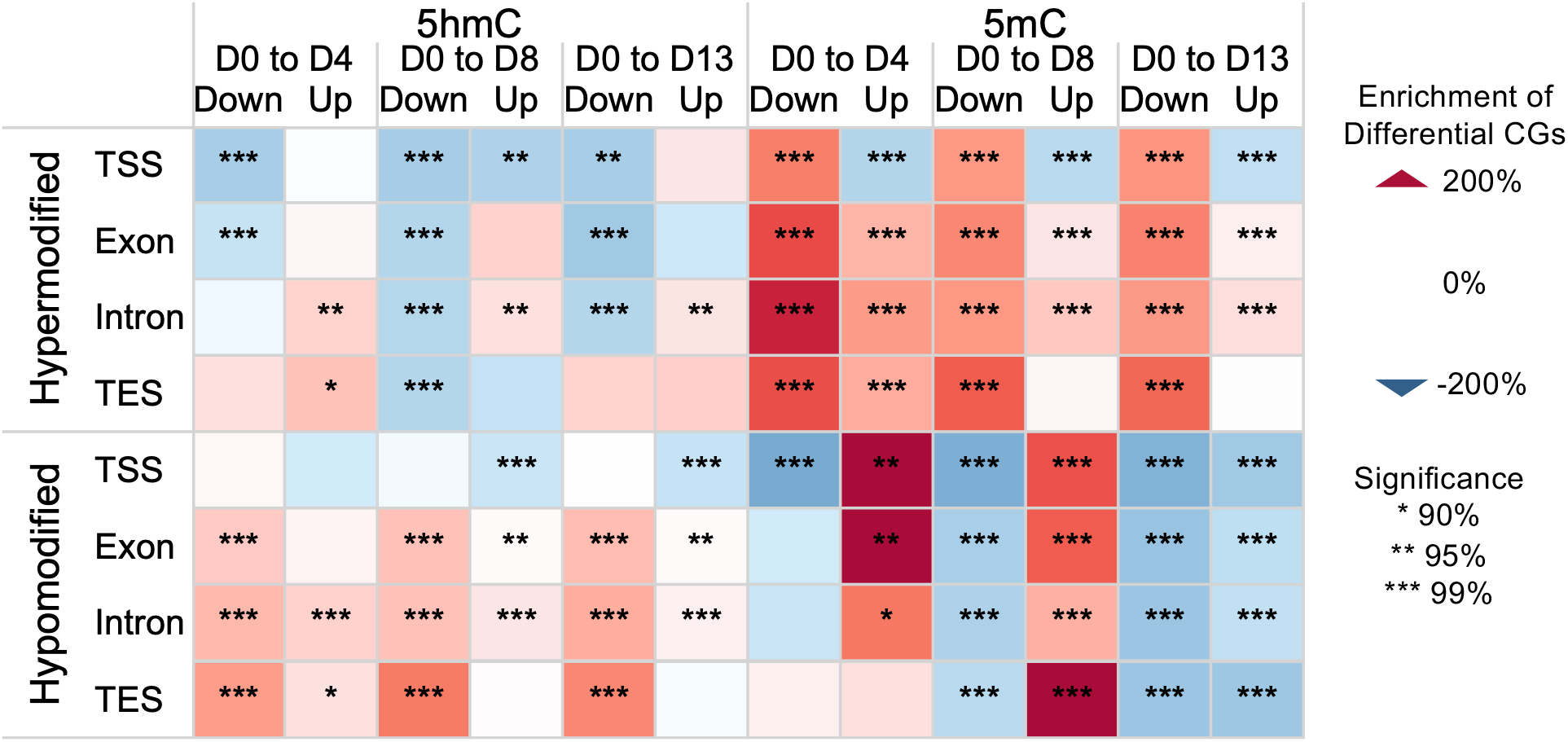
Enrichment of differential CG sites among upregulated and downregulated transcripts relative to non-differentially expressed transcripts for the same day-to-day comparison. Differential CGs within DMRs and DhMRs intersecting with each feature were counted and normalized to the number of CGs in the feature. A T-test was performed between these values for differentially expressed and non-differentially expressed protein-coding transcripts. Enrichment represents the % difference in differential CGs compared to non-differentially expressed transcripts. Additional feature subtypes are shown in Figure S6.

In contrast to the role of 5mC, 5hmC loss was associated mainly with downregulation of gene expression. Downregulated transcripts from D0 compared to D4, D8 and D13 were significantly associated with hypoDhMRs but not hyperDhMRs. Notably, TES regions were the most enriched of all feature types in hypoDhMRs among downregulated transcripts, suggesting this may be an important region in 5hmC mediated gene regulation. Introns and exons within the gene body were also enriched in hypoDhMRs among downregulated transcripts. The same trend was not seen among TSS regions and first exons, where we found fewer changes in hydroxymethylation in either direction. Beyond the gradual erasure of 5hmC throughout the differentiation as observed in figure 3D, we observed enrichment of hypoDhMRs in the gene body and TES relative to non-differentially expressed transcripts. The exception was introns, where we observed hyperDhMRs among upregulated transcripts in all three stages. We also observed a significant association between hypoDhMRs in introns and upregulated transcripts across stages, suggesting loss of 5hmC can be associated with upregulation as well.

### 5-hydroxymethylcytosine to 5-methylcytosine transition regulates key developmental genes

During the ES differentiation process, expression of pluripotent genes gradually decreases, concurrent with increased expression of differentiation marker genes (35). Similarly, during mouse neural development from blastocysts the downregulation of pluripotent genes Oct4, Nanog and Rex1 occurs followed by up-regulation of lineage-specific genes. To understand what role 5mC and 5hmC play in pluripotent gene downregulation, we examined a key neuronal regulator gene, Pax6, along with common representative pluripotent network genes Oct4, Nanog, Rex1 and Rest (Figure 7). Indeed, all four pluripotent representative genes displayed a decrease in 5hmC **with** a concurrent increase of 5mC. As expected, the increase in 5mC levels on these genes resulted in a downregulation of RNA expression. We also observed a common pattern of gain of 5mC on TSS, 1^st^ intron, and gene body suggesting repressive chromatin formation leading to downregulation of the genes (Figure 7B-E). Decreasing 5hmC in the TES region is observed among all four downregulated genes, which is in alignment with the trend observed among downregulated transcripts (Figure 6). Additionally, Nanog, Rex1, and Rest showed 5hmC loss across the gene body, largely overlapping regions of 5mC gain.

**Figure 7.**
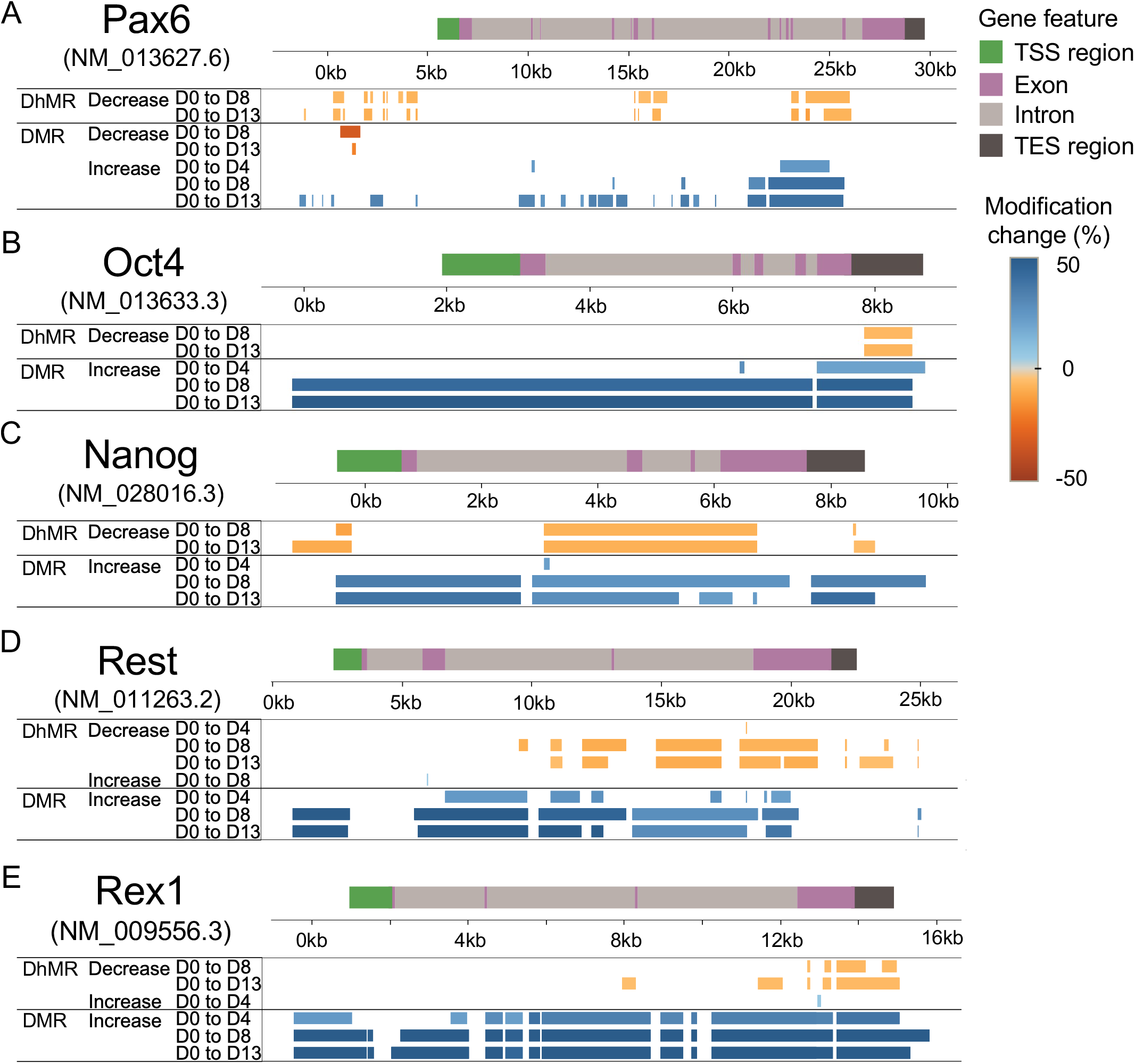
DMR and DhMR locations for differentially expressed transcripts from neuronal gene Pax6. (A), and pluripotent genes Oct4 (B), Nanog (C), Rest (D), and Rex1 (E). The main transcript is shown for each gene, and TSS region, exons, introns, and TES region are indicated. Pax6 (NM_013627.6) was upregulated between D0 and D4 (logFC 7.4, p<0.05). Pluripotent genes were downregulated: Oct4 (NM_013633.3, logFC -11.7 D0 to D13), Nanog (NM_028016.3, logFC -15.5 D0 to D13), Rest (NM_011263.2, logFC -3.4 D0 to D8), and Rex1 (NM_009556.3, logFC -16.8 D0 to D13). DhMRs and DMRs are colored according to change in methylation, with blue indicating hypermethylation and orange indicating hypomethylation. Genomic distance is shown in kb.

Additionally, Pax6, a sequence-specific transcription factor, can activate and repress transcription in multiple cell types in the central nervous system (CNS) demonstrated DNA methylation dynamics. The Pax6 gene has multiple subtypes. And this gene family displayed loss of 5hmC and concurrent gain of 5mC during neuronal differentiation (Figure S7). Among these subtypes, two were downregulated and two were upregulated, suggesting different Pax6 subtypes gain different epigenetic programing of Pax6 during neuronal differentiation (Figure 7A).

## Discussion

Genome-wide mapping of 5hmC has progressed through several generations of technology. Early studies, including our own, relied on enrichment of 5hmC-containing fragments. This was done by immunoprecipitation (hMeDIP-seq), selective chemical labeling, or modification-sensitive restriction digestion (19,36). These approaches established that 5hmC is enriched at enhancers, promoters and gene bodies. However, they report 5hmC enrichment at the fragment scale rather than with single-base resolution. Bisulfite-based methods such as oxBS-seq and TAB-seq provided the first base-resolution 5hmC data (31,37), at the expense of DNA degradation. Bisulfite-free methods, including ACE-seq and CAPS, subsequently removed this limitation, the latter utilizing a complete chemical conversion with a combination of KRuO_4_ oxidation and borane reduction (33,38,39).

More recently, techniques have moved toward single-cell and multiomic measurements. Joint-snhmC-seq, SIMPLE-seq, scTAPS/scCAPS+ and schmC-CATCH resolve 5hmC, alone or together with 5mC, at base resolution in individual cells (40-43). Mx-TOP combines 5hmC and unmodified CG mapping with chromatin accessibility in a single library (44).

E5hmC-seq is a variation of EM-seq in that it shares the T4-BGT and APOBEC chemistry of EM-seq, but without TET2 oxidation (Fig. 1A). EM-seq and E5hmC-seq libraries can therefore be prepared in parallel from the same DNA, and 5mC can be inferred by subtraction. While methods like SIMPLE-seq (42) can profile 5mC and 5hmC from the same molecule, this method utilizes multi-stage chemical and enzymatic conversion, leading to a higher degree of experimental complexity. Moreover, in comparison to the single cell methods described above, which efficiently capture cellular heterogeneity at the expense of sparse CG coverage, E5hmC-seq enables 5hmC analysis genome-wide.

Comparison with hMeDIP-seq in human brain DNA shows that E5hmC-seq offers increased sensitivity compared with enrichment-based methods. In hMe-DIP-seq enrichment is biased toward regions densely modified with 5hmC. While we observed a low false positive rate with hMeDIP enrichment, and the fraction of CGs falling within enriched regions increased with 5hmC level (Figure S1L, M), sensitivity was limited. Of the 7.6 million CGs carrying more than 40% 5hmC, 62% were not significantly enriched by hMeDIP, and only ∼15% of CGs with less than 10% of 5hmC were captured. E5hmC-seq allows for a more complete and precise assessment of 5hmC even at isolated or lower-hydroxymethylation sites.

Our results demonstrate that E5hmC-seq provides robust detection of 5hmC across a broad range of DNA inputs and 5hmC abundances relevant to biological samples. Despite the relatively low 5hmC level in E14 ESCs used here, single-base 5hmC measurements were correlated across DNA inputs from 200 ng to 0.1 ng. Correlations improved when 5hmC levels were aggregated across 1 kb genomic windows. This consistency was demonstrated using genomic windows one-tenth the size of those used in previous studies. E5hmC-seq also successfully reflected the diversity of endogenous 5hmC abundance across 14 mouse tissues. These measurements were also consistent with independent LC-MS/MS quantification, further supporting the accuracy and robustness of this method.

During the ES differentiation to embryoid body formation, the 5hmC decrease with concurrent increase in 5mC reflected the dynamics and balance between both marks. A similar relationship was previously reported during ES differentiation using hMeDIP-seq, where loss of 5hmC and gain of 5mC at promoters accompanied gene silencing (36). Additionally, we show that, during neuronal progenitor cell generation and differentiation, the levels of both 5mC and 5hmC remained at a steady-state level, although transcriptional output was dynamic. The differentially expressed transcripts (both up- and down-regulated) increased from 17K to 24K between D4 to D13, concurrent with dynamic shuffling of DhMRs and DMRs. The dynamic shuffling falls into two categories (1) decrease in DhMRs that had no overlap with DMRs demonstrating their erasure. (2) decrease in DhMRs that had overlap with DMRs with an increase in overall DMRs suggesting erasure of 5hmC followed by gain of 5mC in the genome perhaps. This phenomenon demonstrates rapid epigenetic mark erasure and re-establishment during ES differentiation to neuronal cells.

To relate epigenetic changes to transcript expression among specific genic features, we counted differentially modified CGs within feature types (TSS, exons, introns, and TES). We normalized these counts to the CG content of each feature and compared differentially expressed transcripts against non-differentially expressed transcripts for each comparison to test whether certain feature types were enriched in 5mC and 5hmC changes. This helped us distinguish changes in 5mC and 5hmC relative to the genome-wide erasure of 5hmC that occurred throughout differentiation.

We observed the canonical model of 5mC regulation whereby hyperDMRs were enriched in all feature types examined among downregulated transcripts at D4, D8, and D13. HypoDMRs at the TSS and within gene bodies were enriched among upregulated transcripts through NPC generation (D8), possibly suggesting that loss of 5mC contributes to early differentiation.

5hmC showed an asymmetric relationship whereby the loss of 5hmC was enriched among downregulated transcripts during all three stages. Both 5mC gain and 5hmC loss are correlated with downregulation during differentiation, but the 5hmC observation was strongest at the TES, followed by introns and exons. Gain of 5hmC did not show a similar association with upregulation, however far fewer hyperDhMRs were observed at each stage, so our power to observe these relationships was limited. The one notable exception to this was among introns, where we did observe a significant relationship between upregulated transcripts and gain of 5hmC during all three stages. The pluripotency genes examined (Oct4, Nanog, Rex1, Rest) each showed loss of 5hmC at the TES (Figure 7), which is consistent with the transcriptome-wide observations. Aside from Oct4, these genes showed patterns of 5hmC loss to varying degrees throughout the gene body, generally overlapping regions of 5mC increase.

These results suggest that during neuronal differentiation, 5hmC changes within gene bodies and near the TES are important marks of transcriptional regulation. This agrees with the prior association of 5hmC within the gene body associated with active transcription in neural cells (45) and with recent data implicating gene-body 5hmC dynamics in neuronal differentiation (44). We extend these observations by identifying 5hmC loss near the TES as a prominent marker of gene silencing.

In summary, profiling of 5mC and 5hmC across E14 differentiation revealed progressive erasure of the 5hmC landscape during embryoid body formation, NPC generation, and neuronal differentiation. Approximately half of the regions that lost 5hmC concurrently gained 5mC. These changes are concentrated within gene bodies and towards the TES and correlated with downregulation of pluripotent genes. Together, these findings highlight gene-body 5hmC, and not just promoters, as a dynamic feature of developmental gene regulation and underscores the necessity of single-base 5hmC analysis for understanding of epigenetic regulation.

## Supplemental Figures

**Figure S1.**
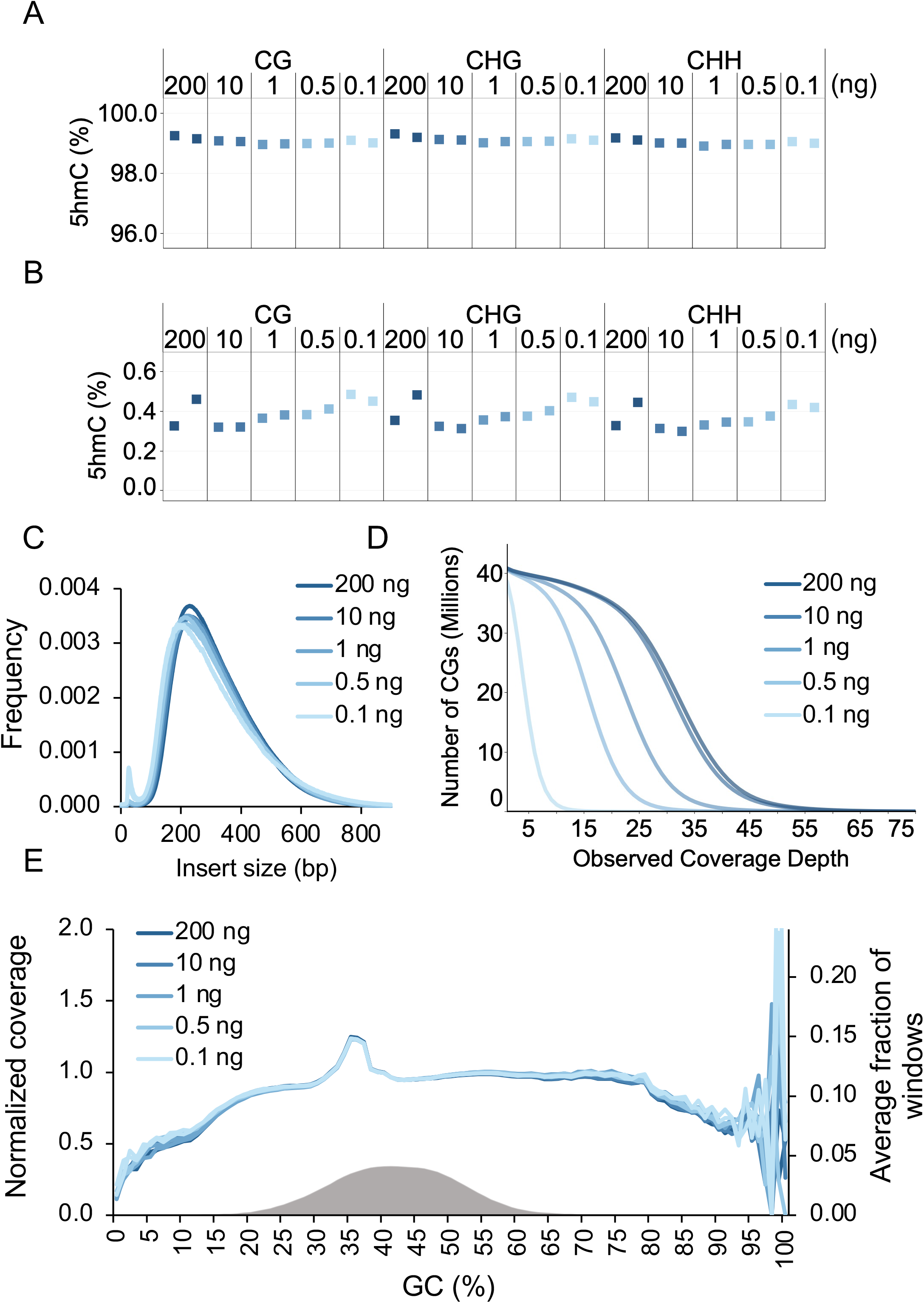
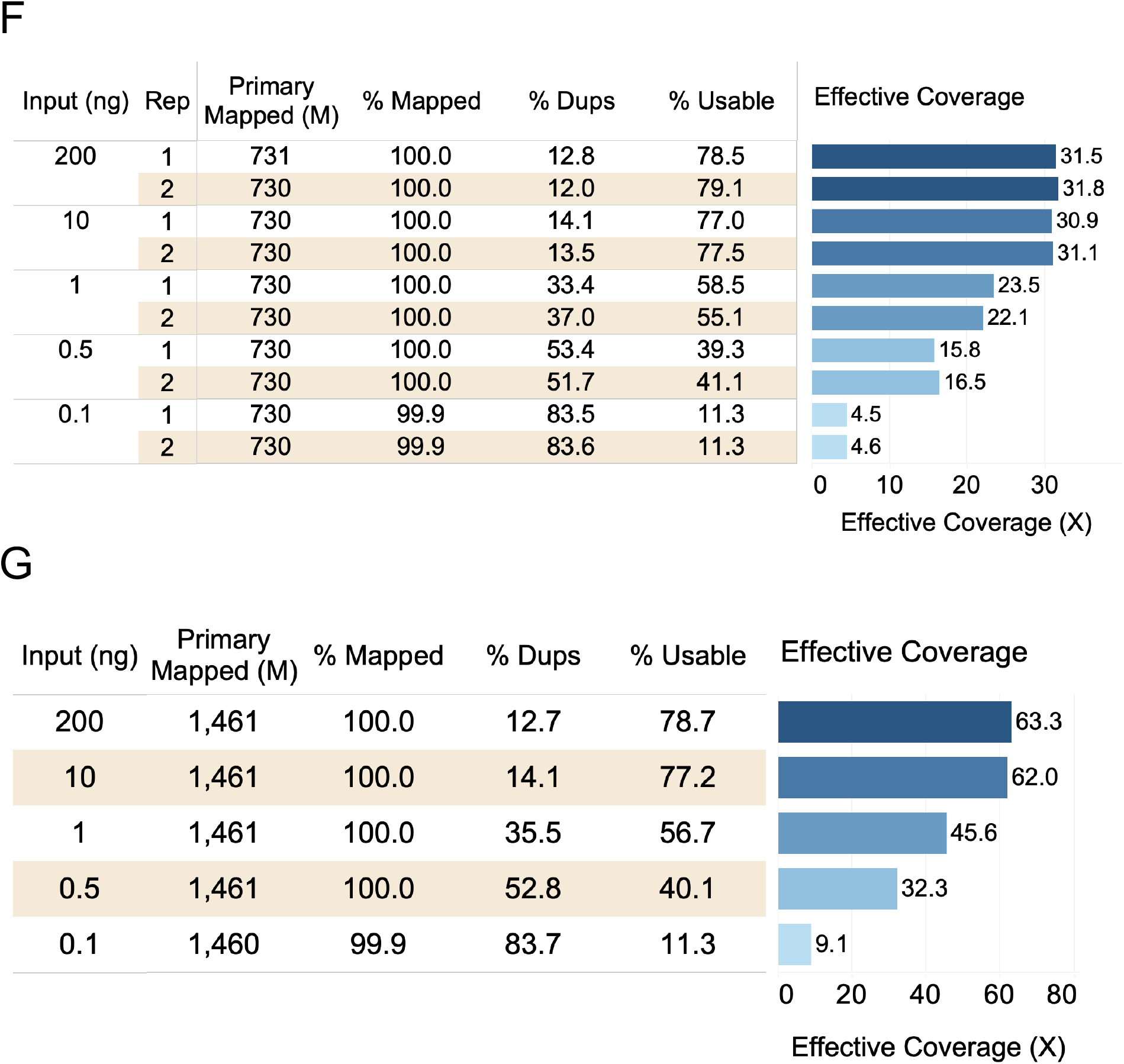
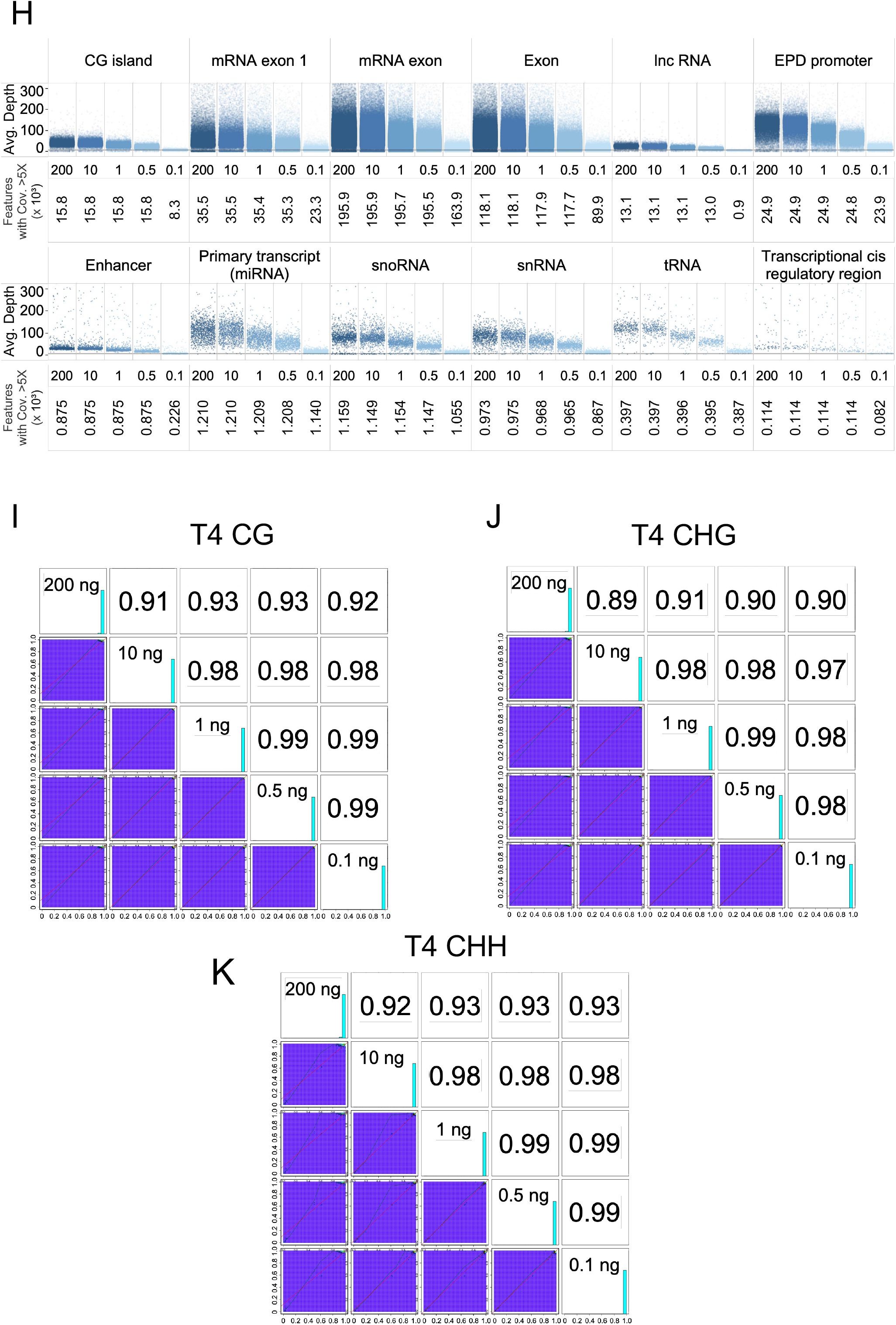
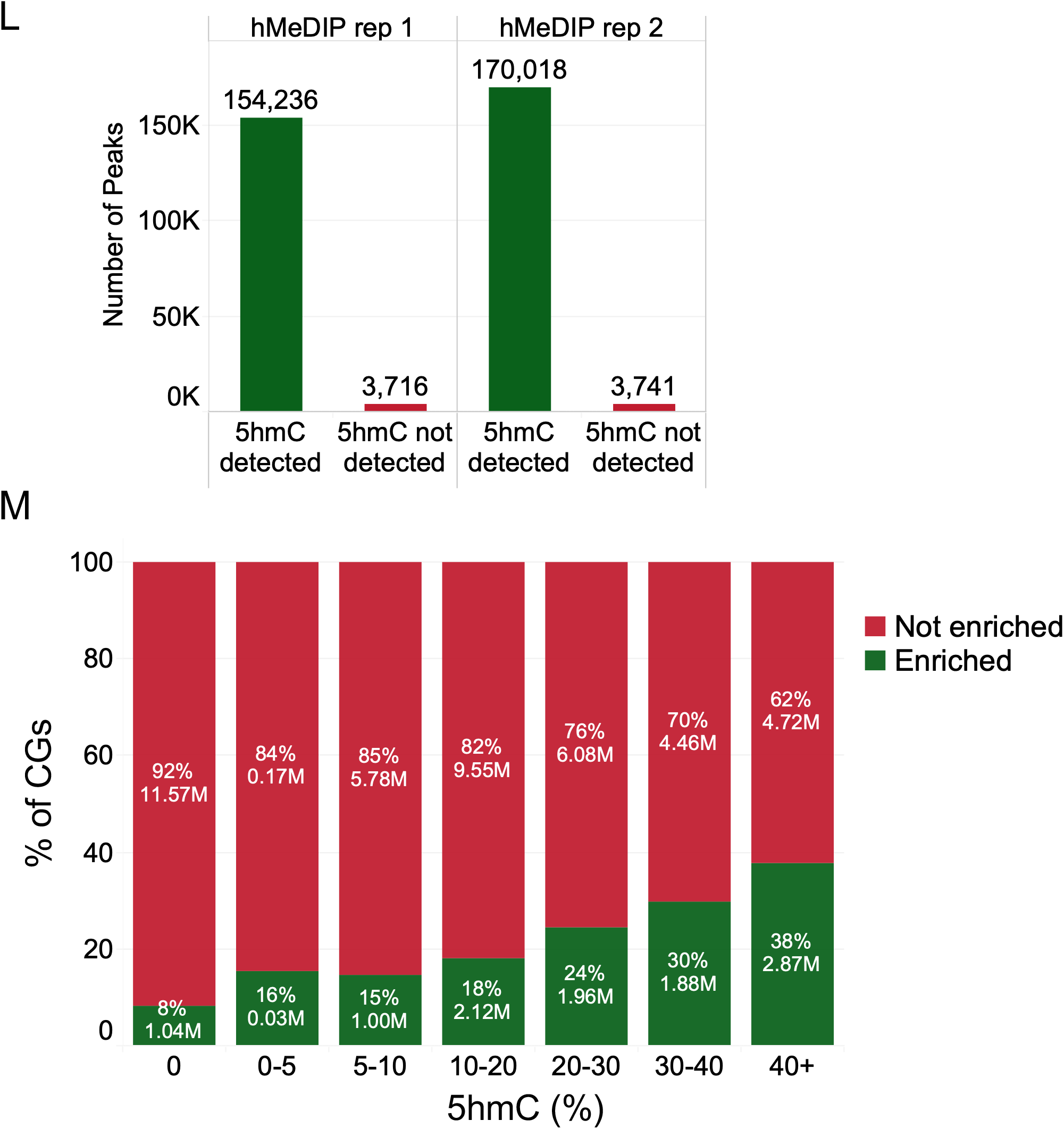
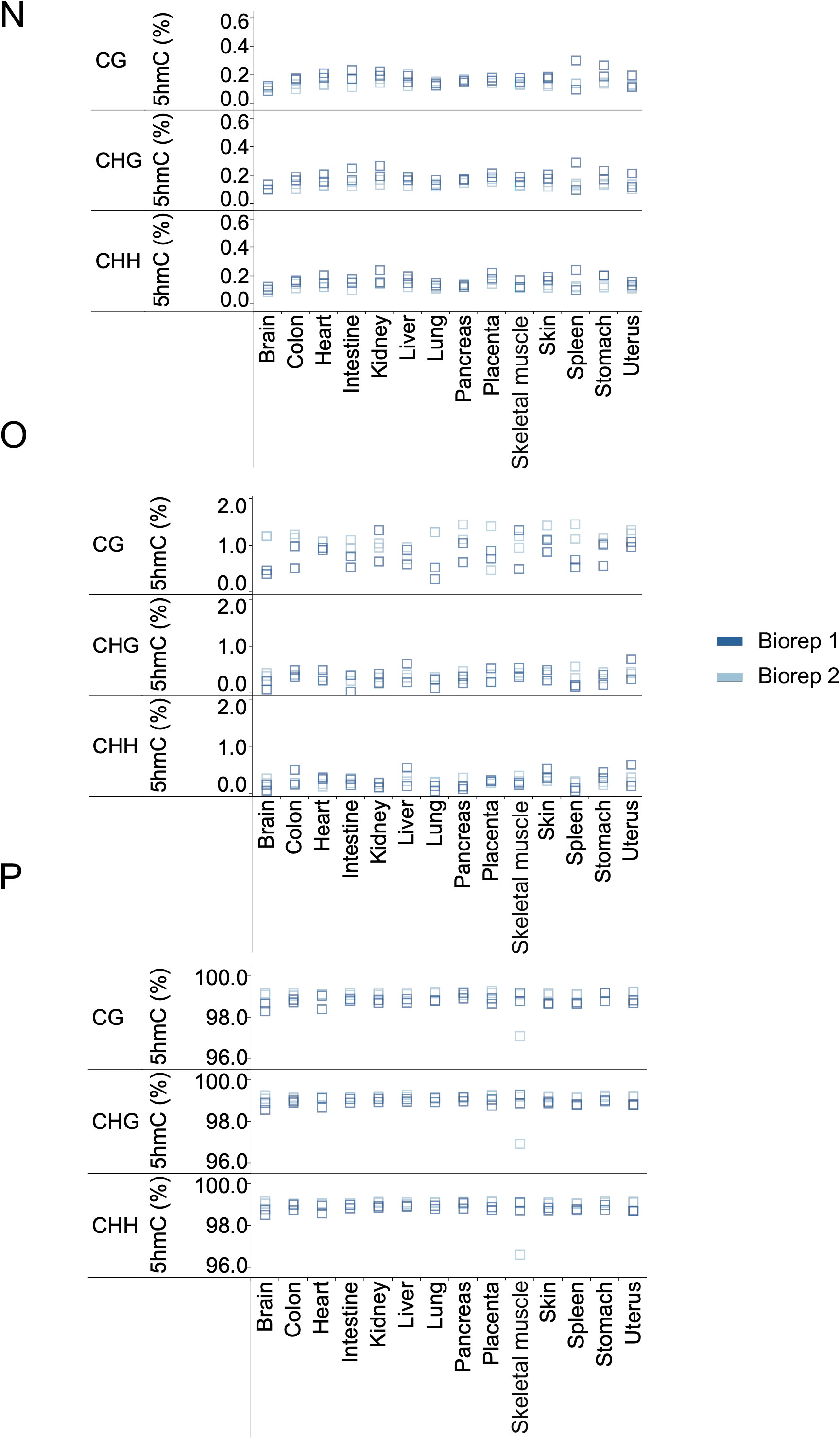
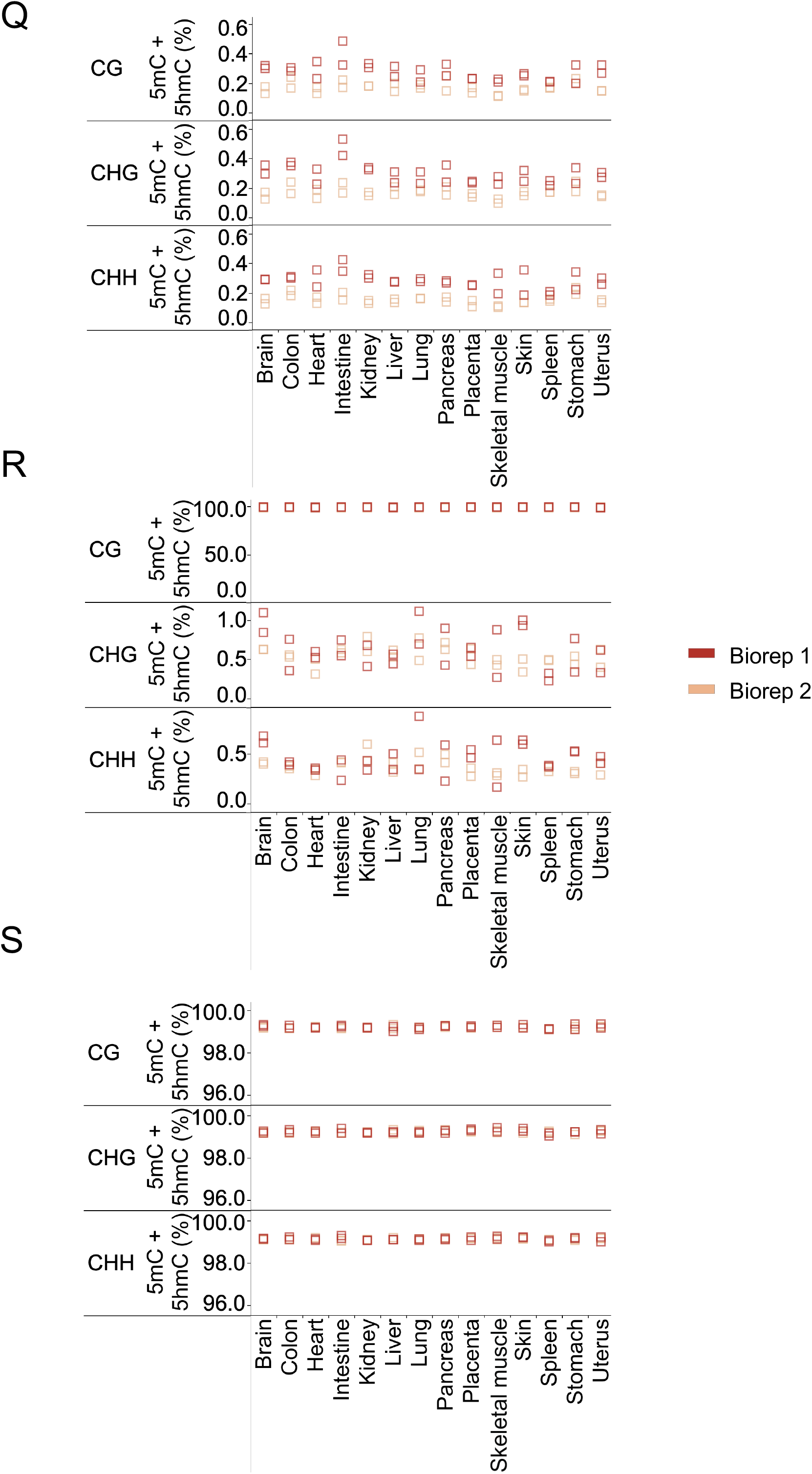
Qualitative analysis of E5hmC-seq. (A) Percent 5hmC level detected in fully hydroxymethylated T4 DNA in CG, CHG, and CHH using E5hmC-seq libraries. (B) Percent 5hmC level detected in fully unmethylated lambda phage DNA in CG, CHG and CHH with E5hmC-seq libraries. (C) Insert size distribution of E5hmC-seq libraries with various amounts of E14 gDNA. (D) Cumulative coverage plot showing CG coverage at 1X – 75X sequencing depth for different input gDNA libraries. The 200 ng – 0.5 ng inputs detected at least 40 million CGs at 1X depth and the 0.1 ng input detected ∼38.7 million CGs at 1X depth. (E) GC coverage in E5hmC-seq libraries was analyzed using Picard and the distribution of normalized coverage across different GC contents of the genome (0-100%) was plotted. Distribution of GC content in the mouse genome is shown in grey. Note that the higher region in between 30 and 40% GC is a result of a copy-number mismatch between the reference genome and the sample. (F) Sequencing and alignment metrics for E5hmC-seq libraries using 730 million total reads. Metrics were calculated using bwa-meth, SAMtools, and Picard. Theoretical coverage is calculated using the number of bases sequenced/total bases in the GRCm39 reference. (% Mapped) Reads aligned to the reference genome (GRCm39+controls); (% Dups) reads marked as duplicate by Picard MarkDuplicates; (% Usable) the set of Proper-pair, MapQ 10+, primary, nonduplicate reads used in methylation calling (SAMtools view -F 0xF00 -q 10); and (Effective Coverage) %Usable x theoretical coverage. (G) Same as (F) except after combining reads for the two technical replicates of each input amount. (H) Base coverage of diverse genomic feature types across inputs. The number of features with coverage greater than 5X is indicated below for each plot. Coverage of genomic feature types is represented with one point per region with the vertical position representing the average coverage of the feature. Points are staggered horizontally to avoid excess overlapping. CG islands were defined based on the UCSC genome browser, and EPD promoters are from the Eukaryotic Promoter Database (EPD). The other feature annotations are from NCBI RefSeq (GCF_000001635.27 GRCm39). Pearson correlations for CG (I), CHG (J) and CHH (K) sites in the fully hydroxymethylated T4 spike-in control. Data shown for each input is the combination of two technical replicate libraries. Correlations were generated at a single base level for 200 ng – 0.1ng at 1X minimum coverage. (L) Peaks were called on two reps of 1ng UltraII hMeDIP libraries using the input control. The number of peaks where 5hmC was detected in deep-sequenced high-input (200ng) human brain 5hmC-seq data (at least one site greater than 0%) is shown. (M) Enrichment of CGs in human brain hMeDIP relative to 5hmC levels in deep-sequenced high-input (200ng) human brain 5hmC-seq libraries. Peaks were called on two reps of 1ng UltraII hMeDIP libraries using the input control. 53M CGs with at least 5X observed coverage in 5hmC-seq data were binned by 5hmC modification level and checked for overlap with peaks. CGs that fell within peaks were considered significantly enriched. The average for the two replicates is shown. Percent CG, CHG and CHH 5hmC detected in (N) unmethylated lambda, (O) CG methylated pUC19 and (P) fully hydroxymethylated T4 for mouse tissue survey E5hmC-seq libraries. Similarly, percent CG, CHG and CHH methylation (5mC + 5hmC) detected (Q) unmethylated lambda, (R) CG methylated pUC19 and (S) fully hydroxymethylated T4 for mouse tissue survey EM-seq libraries.

**Figure S2.**
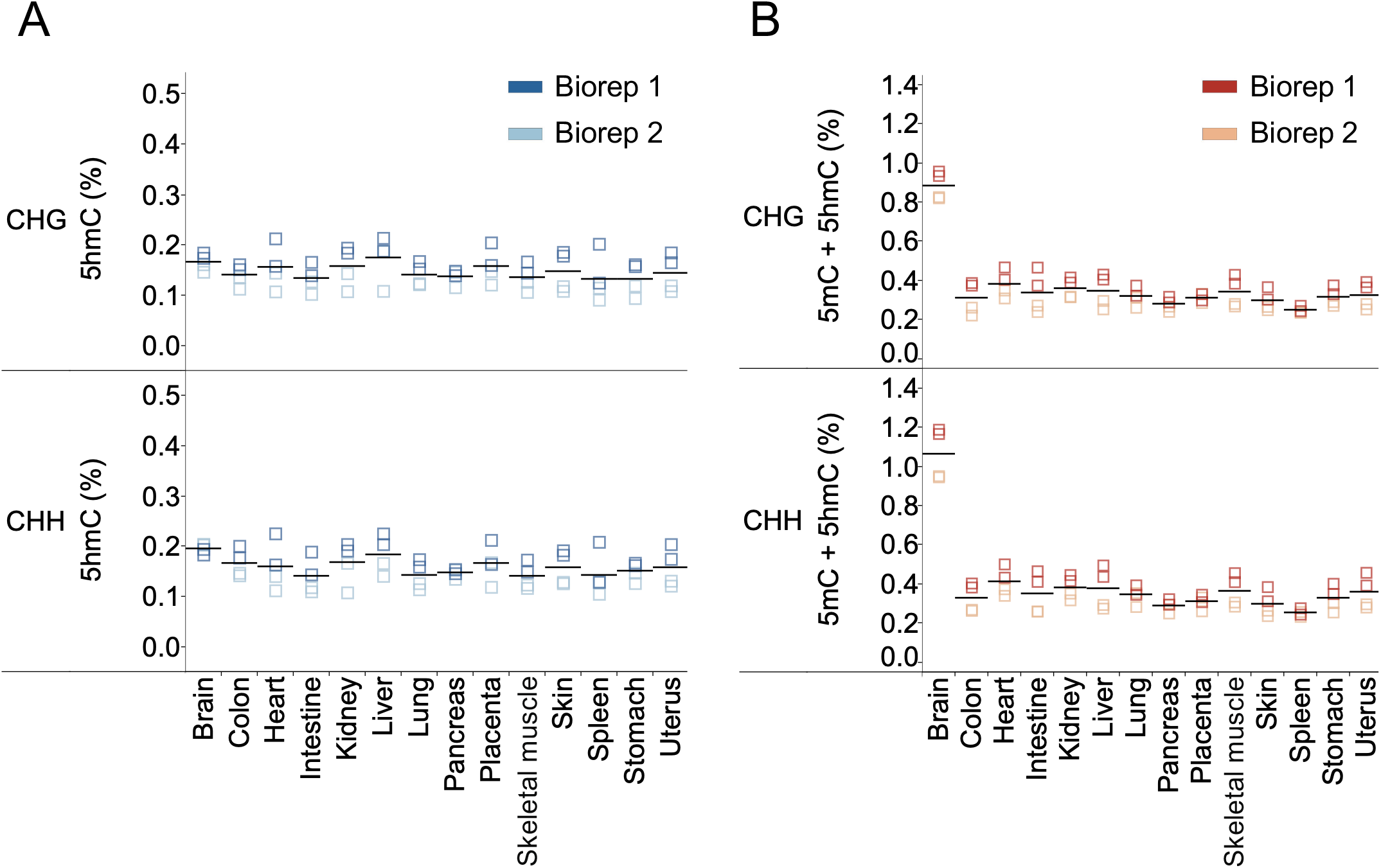
Methylation profiling in various mouse tissue gDNA (A) Percent CHG and CHH 5hmC detected in E5hmC-seq libraries. (B) Percent CHG and CHH methylation (5mC + 5hmC) in the same mouse gDNA from EM-seq data.

**Figure S3.**
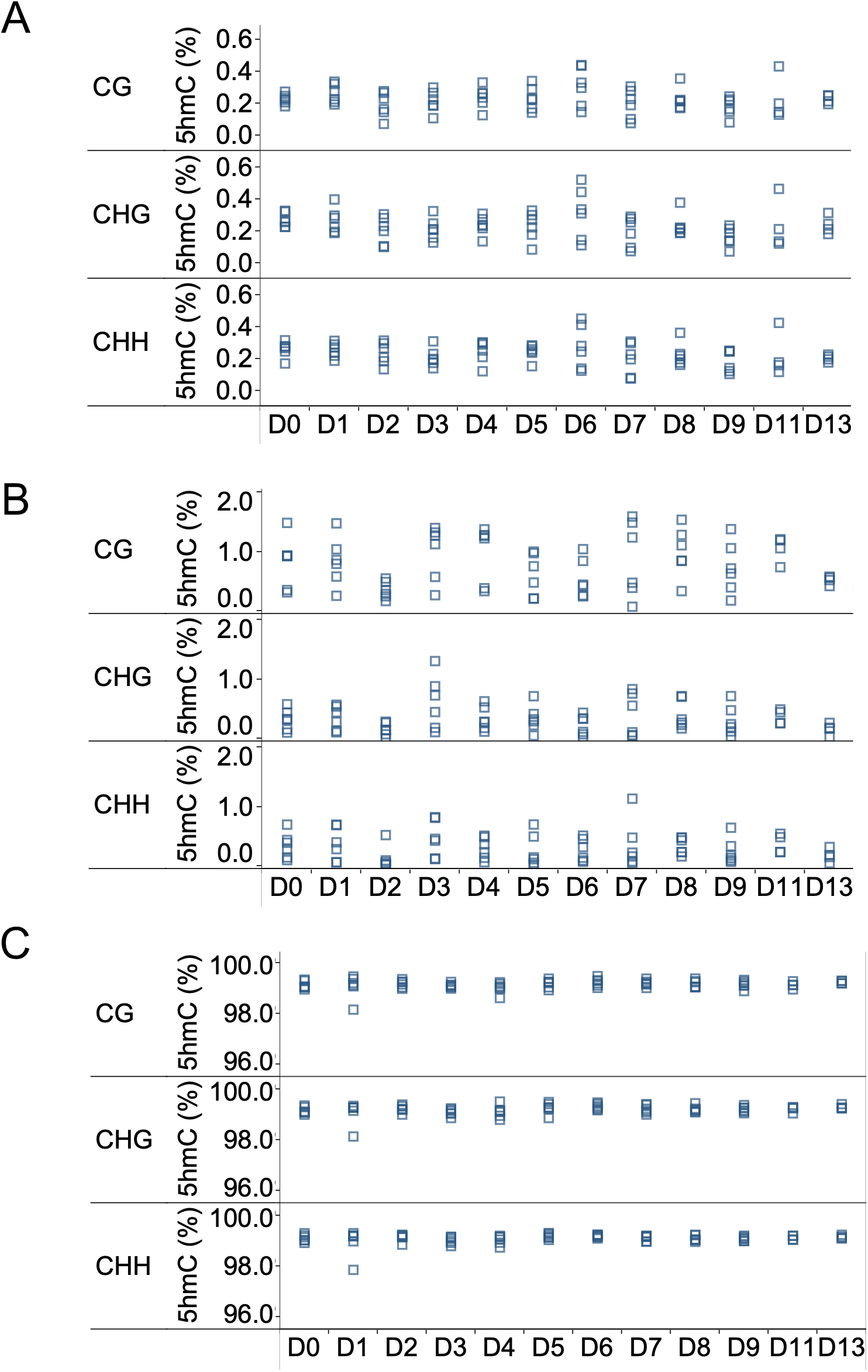
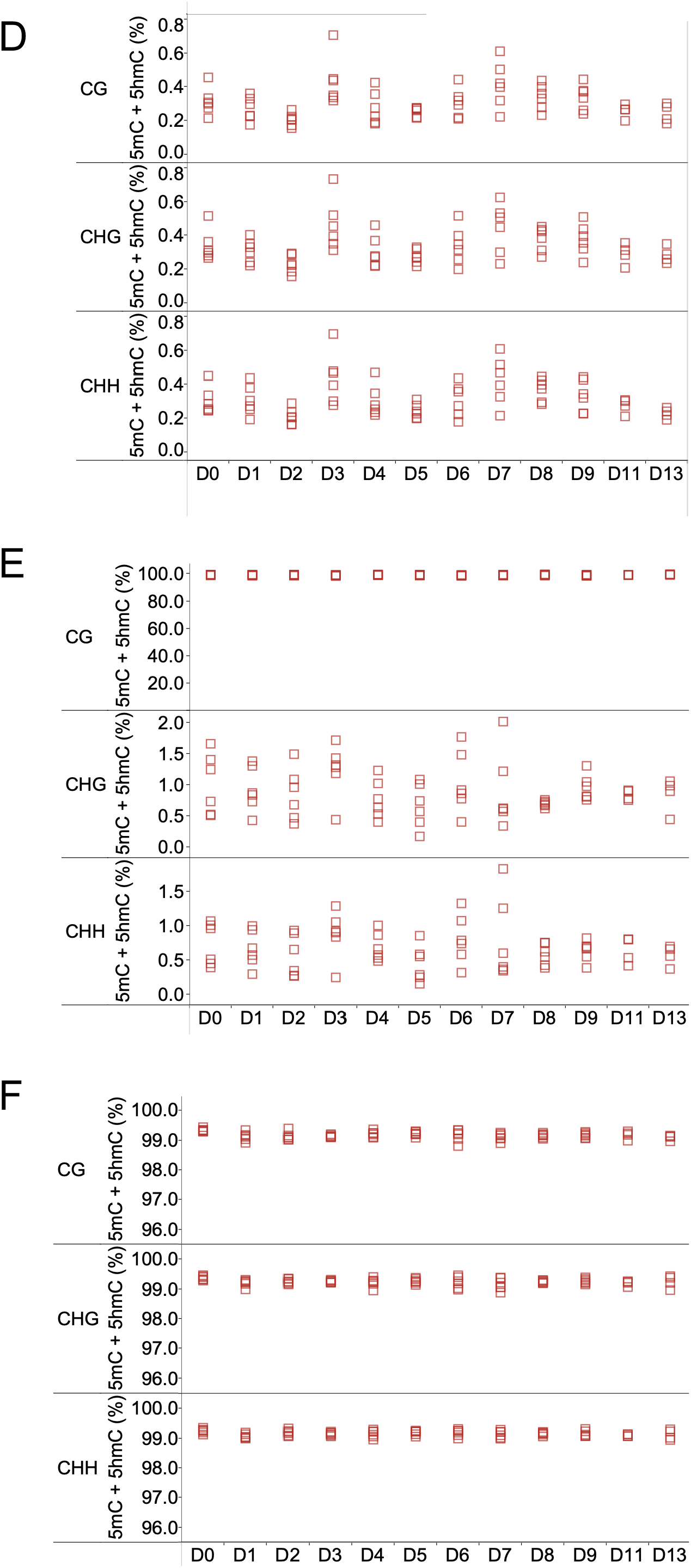
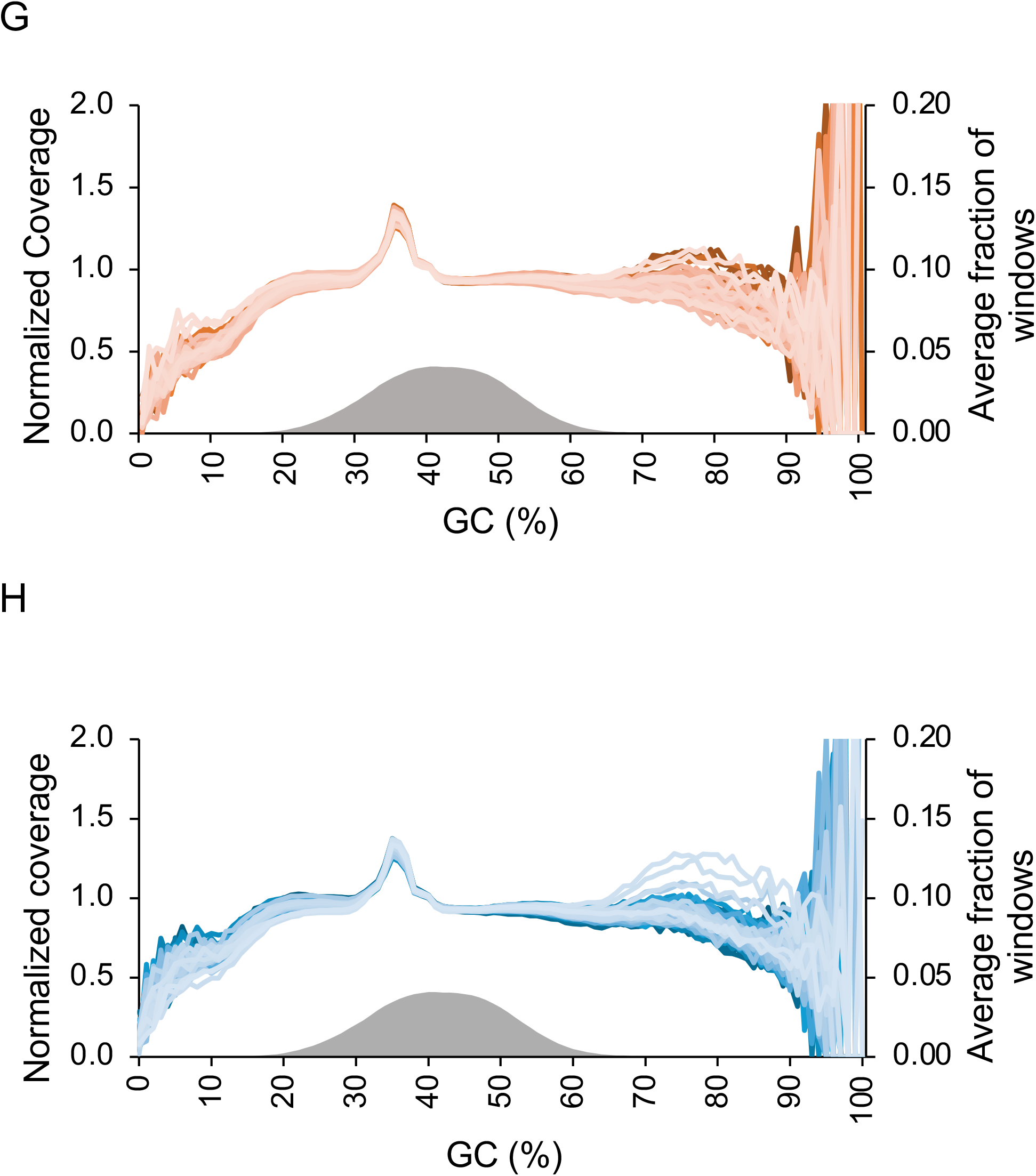
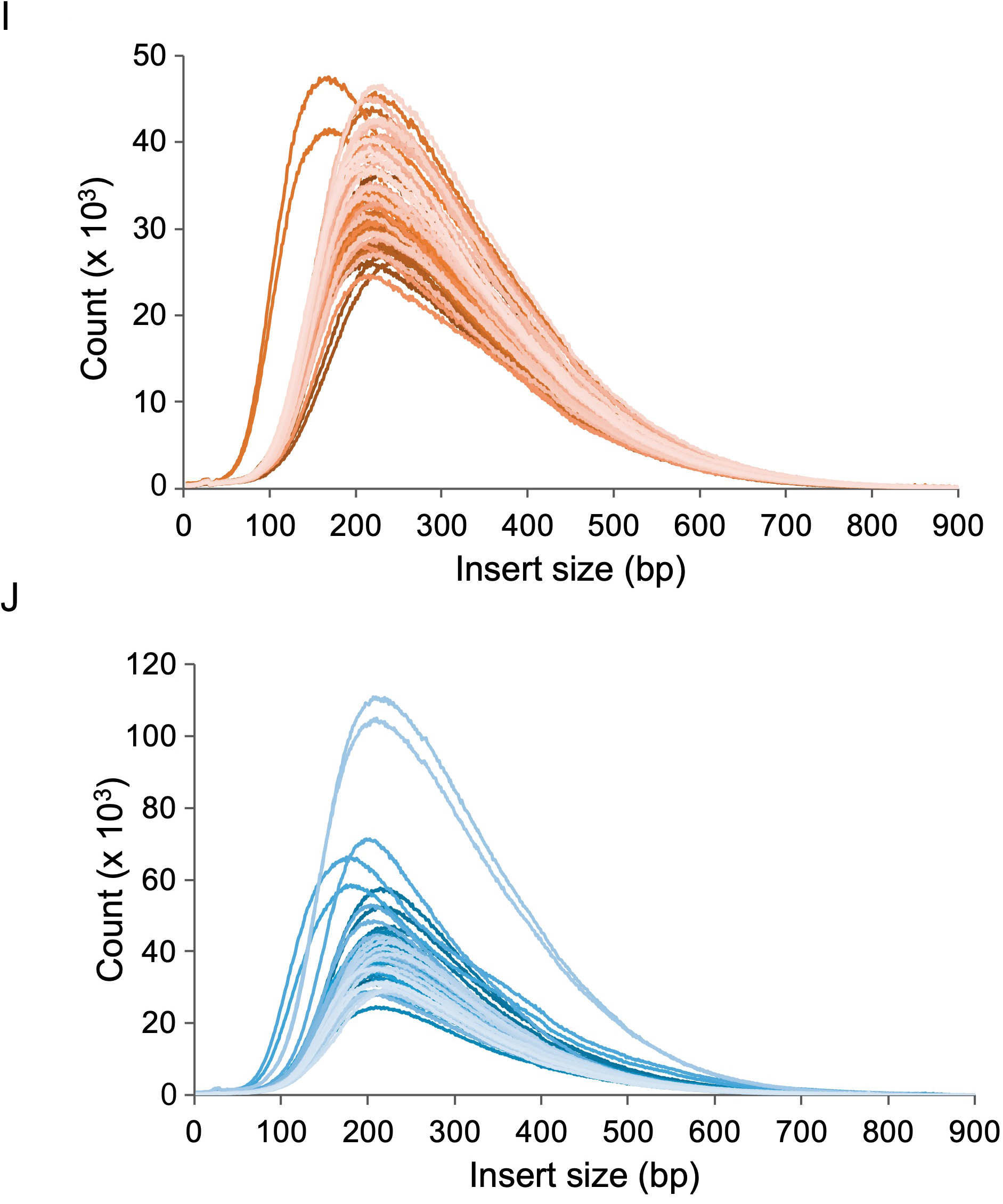
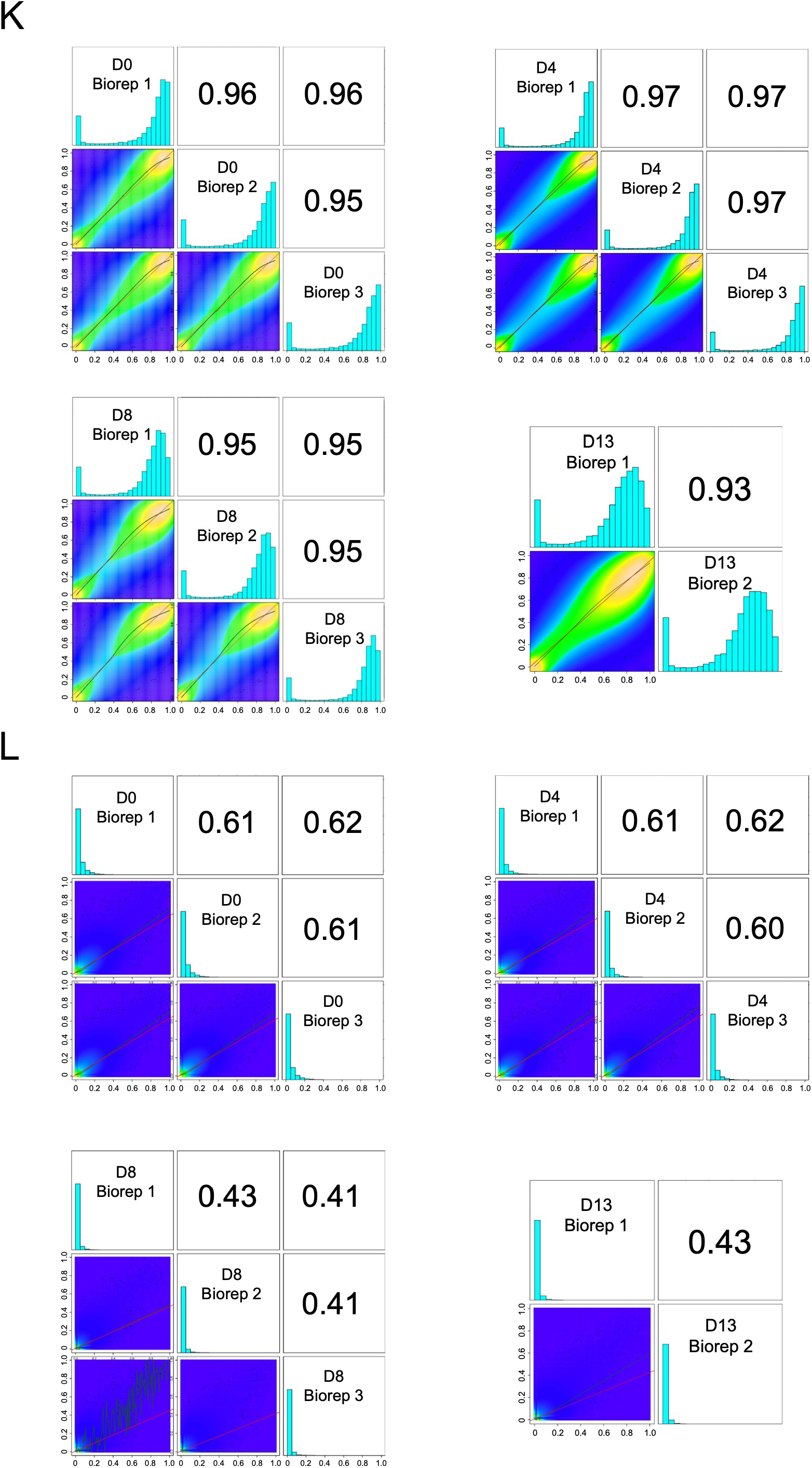
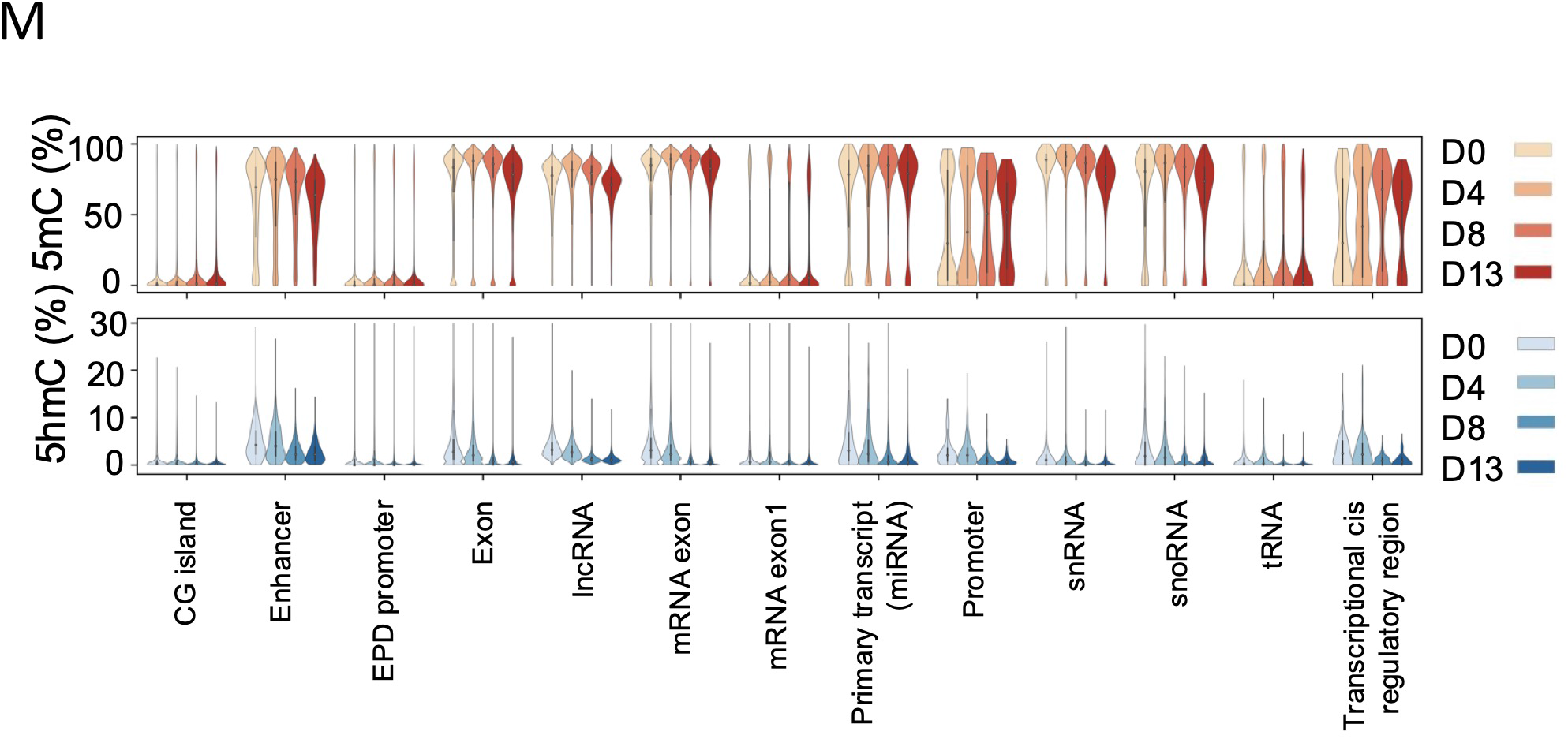
Qualitative analysis of E5hmC-seq for E14 differentiation time course. Percent CG, CHG and CHH 5hmC detected (A) unmethylated lambda, (B) CG methylated pUC19 and (C) fully hydroxymethylated T4 for E14 differentiation time course E5hmC-seq libraries. Percent CG, CHG and CHH methylation (5mC + 5hmC) detected in (D) unmethylated lambda, (E) CG methylated pUC19 and (F) fully hydroxymethylated T4 for E14 differentiation time course EM-seq libraries. GC coverage in (G) EM-seq or (H) E5hmC-seq E14 differentiation libraries were analyzed using Picard and the distribution of normalized coverage across different GC contents of the genome (0-100%) was plotted. Distribution of GC content in the mouse genome is shown in grey. Note that the higher region in between 30 and 40% GC is a result of a copy-number mismatch between the reference genome and the sample. Insert size distribution of (I) EM-seq or (J) E5hmC-seq of E14 differentiation libraries. Pearson correlations between individual bioreps of EM-seq (K) or E5hmC-seq (L) E14 differentiation libraries. All correlations are at 1X minimum coverage. Data shown for each library is the combination of two technical replicates. (M) Distribution of average 5mC and 5hmC methylation level across feature types.

**Figure S4.**
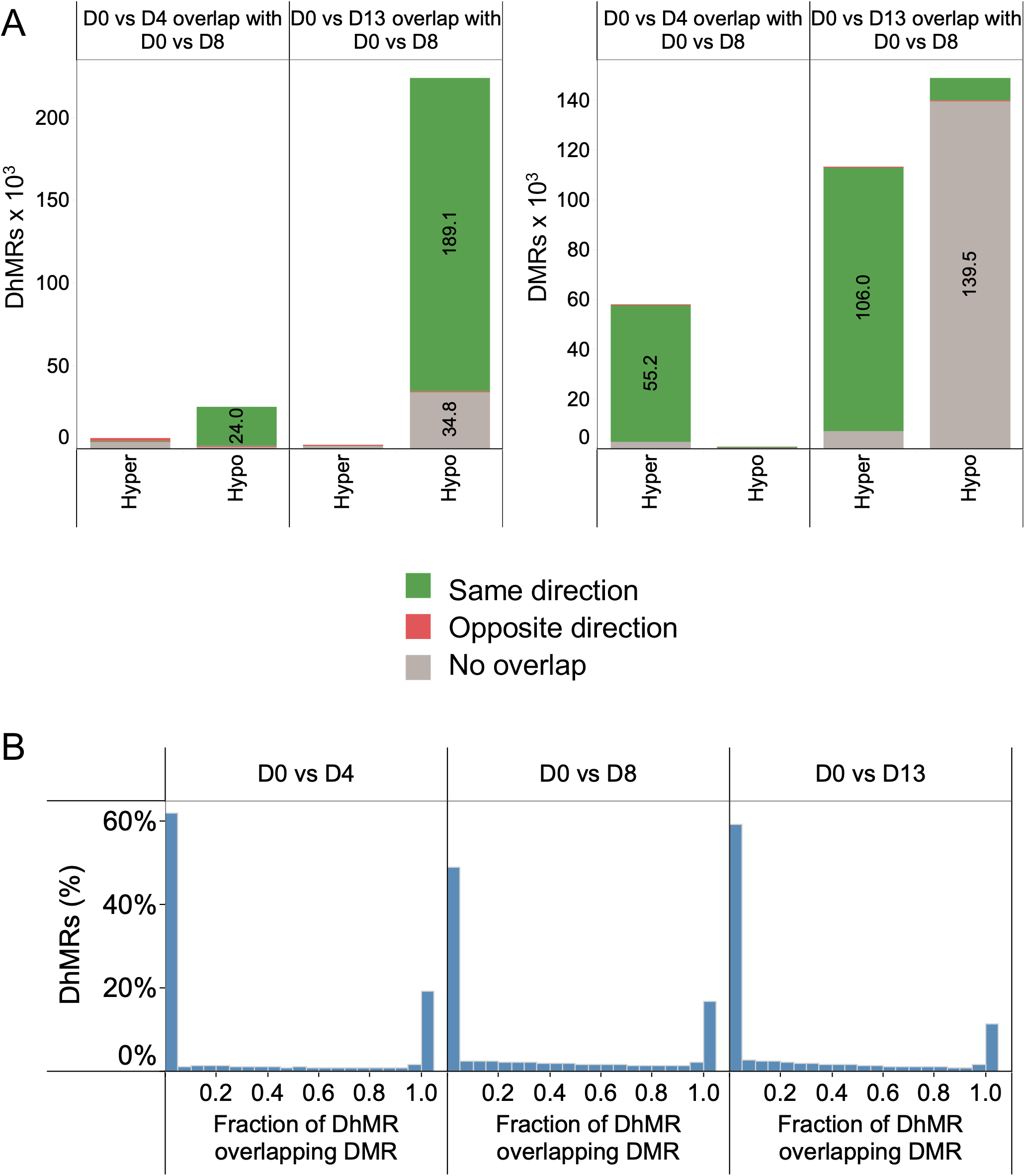
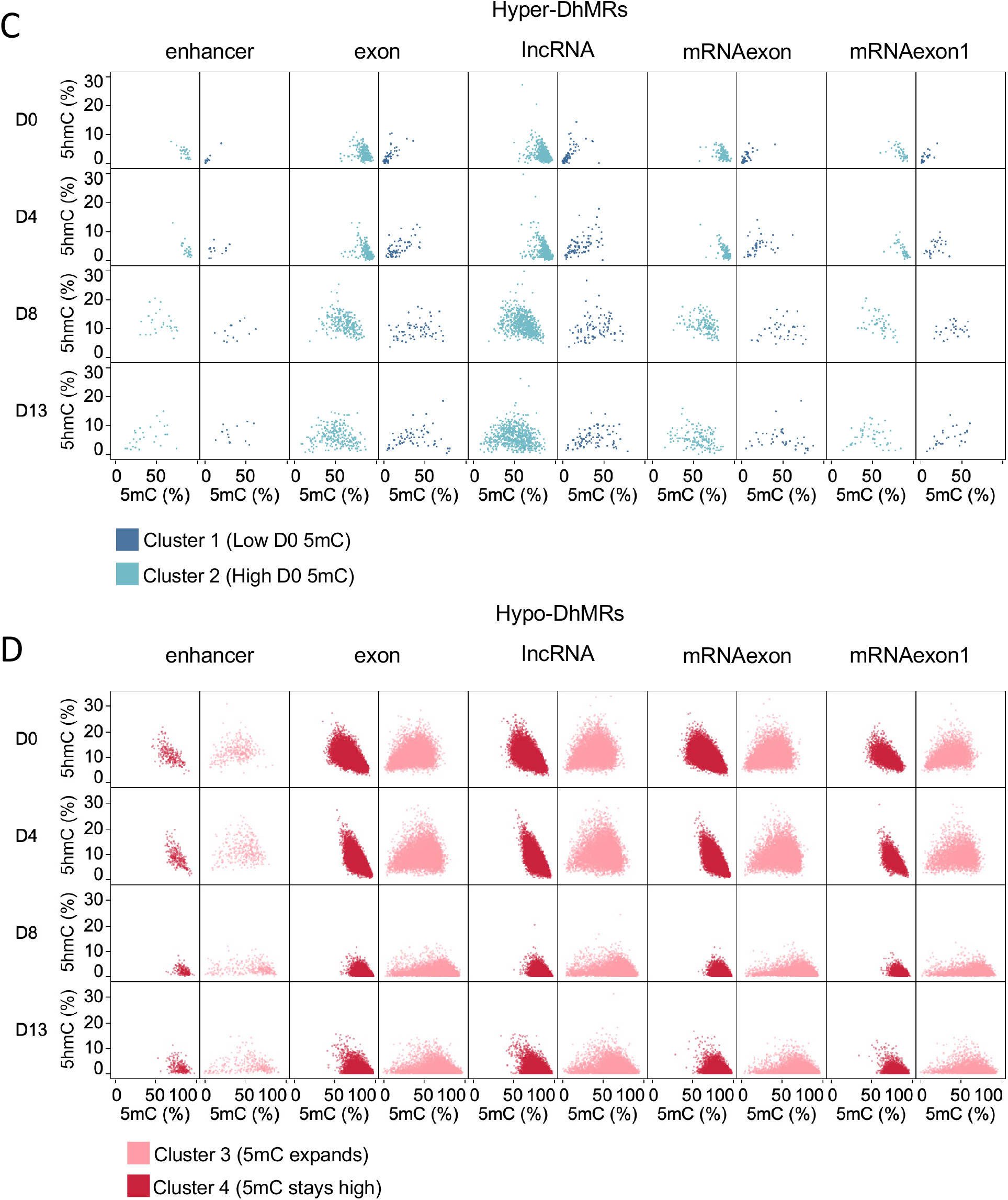
Differential methylation and hydroxymethylation during E14 differentiation. (A) The overlap of D0 vs D4 and D0 vs D13 DhMRs (left) and DMRs (right) with D0 vs D8, was calculated requiring a 20% overlap for either region. Overlapping regions were further categorized into those with the same direction of change (green) and those with opposite directions of change (red). (B) Distribution of proportion of DhMRs overlapping DMRs. DhMRs were intersected with DMRs from the same day comparison (e.g. D0 vs D4 DhMRs overlapping D0 vs D4 DMRs). The length of the overlap was divided by the length of the DhMR. Mean 5hmC and 5mC methylation levels of hyper-DhMRs (C) and hypo-DhMRs (D) from the D0 vs D8 comparison intersecting various feature types. A 1bp overlap was required between the DhMR and the feature type. The same clusters from Figure 4 are shown.

**Figure S5.**
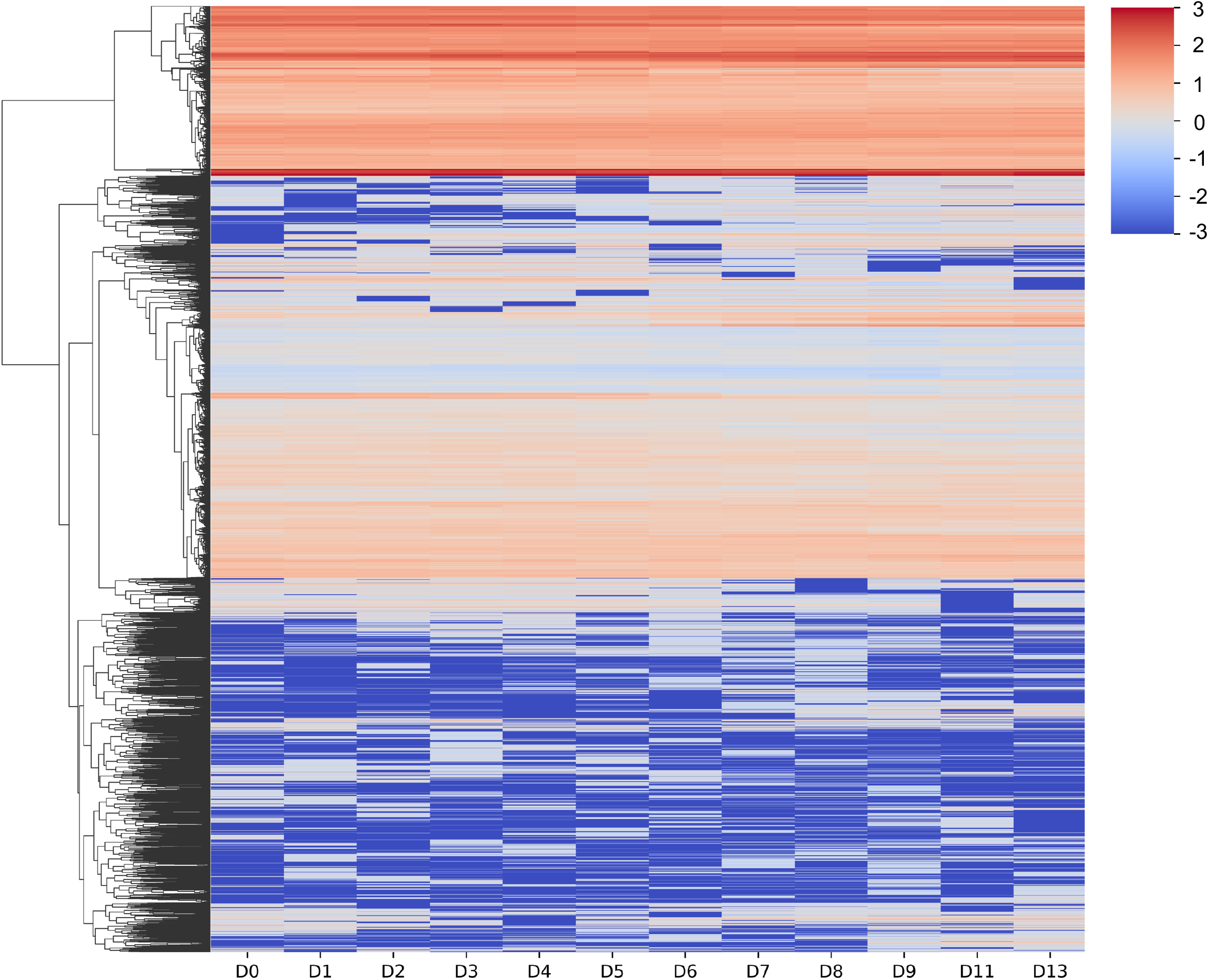
Heatmap displaying pattern of TPM transition from D0 to D13 time points (within a power transformed range ±3)

**Figure S6.**
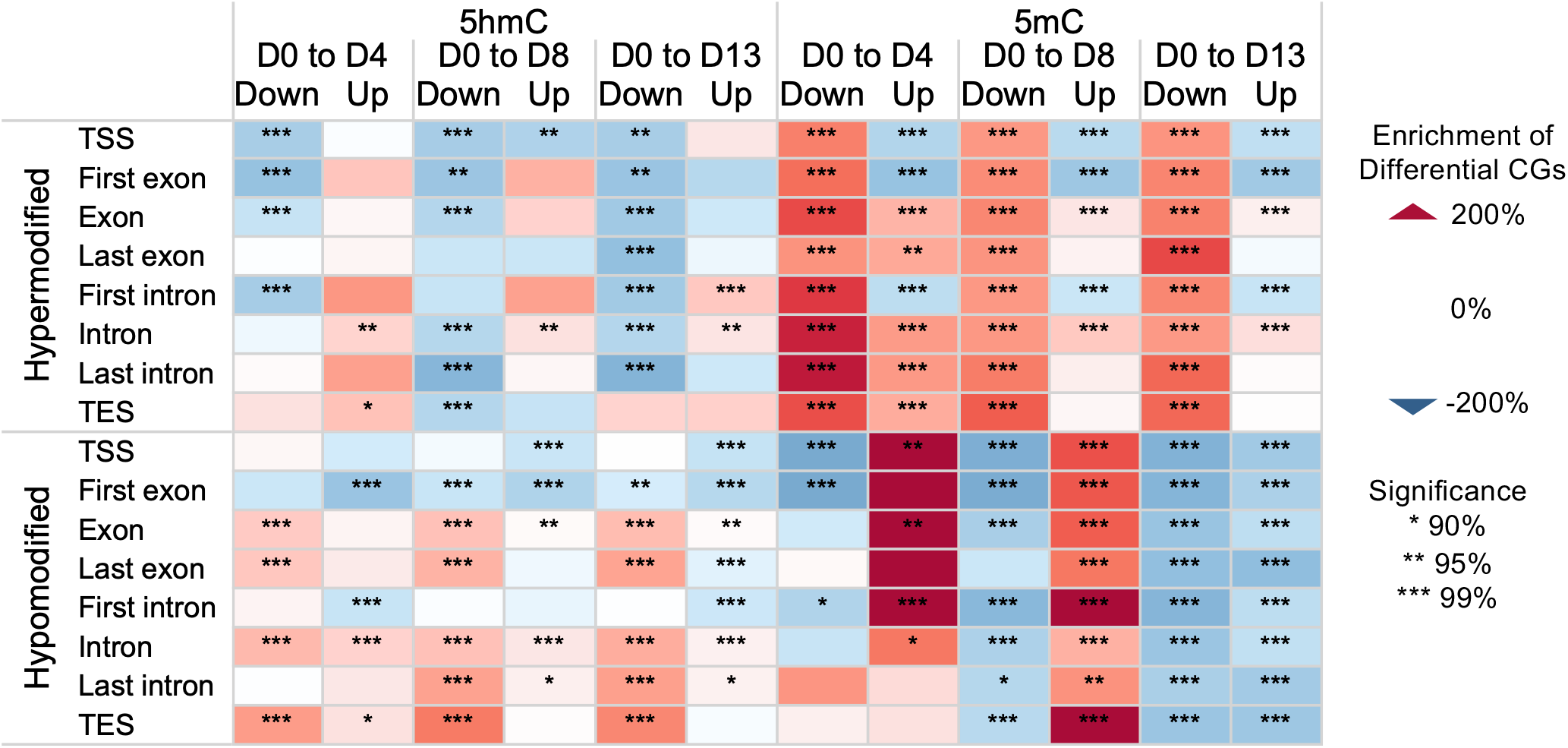
Enrichment of differential CG sites among upregulated and downregulated transcripts relative to non-differentially expressed transcripts for the same day-to-day comparison. Additional categories are shown along with those in Figure 6: first exon, last exon, first intron, and last intron.

**Figure S7.**
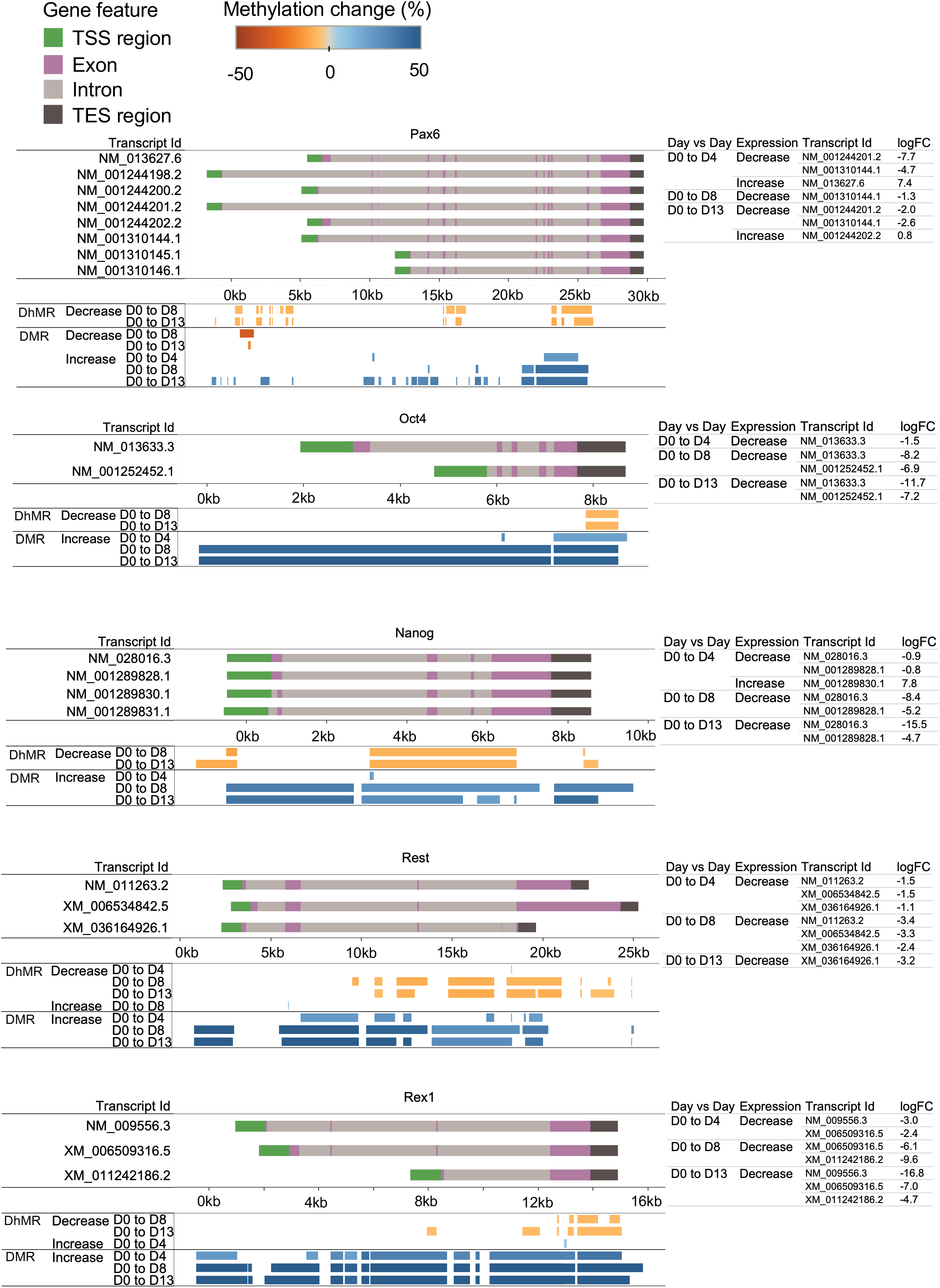
Additional differentially expressed transcripts for genes shown in figure 7, as well as the log fold change values for transcripts that were differentially expressed.

**Supplemental Table S1:** Sequencing and alignment metrics for E5hmC-seq and EM-seq libraries for the mouse tissue survey. Metrics were calculated using bwa-meth, SAMtools, and Picard. (% Mapped) Reads aligned to the reference genome (GRCm39+controls); (% Dups) reads marked as duplicate by Picard MarkDuplicates; (% Usable) the set of Proper-pair, MapQ 10+, primary, nonduplicate reads used in methylation calling (SAMtools view -F 0xF00 -q 10).

| Tissue | Library Type | Biorep | Rep | Primary Mapped (M) | % Mapped | % Dups | % Usable | Tissue | Library Type | Biorep | Rep | Primary Mapped (M) | % Mapped | % Dups | % Usable |
|---|---|---|---|---|---|---|---|---|---|---|---|---|---|---|---|
| Brain | E5hmC-seq | 1 | 1 | 28 | 99.9 | 4.0 | 86.1 | Pancreas | E5hmC-seq | 1 | 1 | 33 | 99.9 | 8.6 | 79.3 |
|  |  |  | 2 | 25 | 99.9 | 3.6 | 86.7 |  |  |  | 2 | 19 | 99.9 | 6.7 | 80.3 |
|  |  | 2 | 1 | 26 | 99.9 | 10.5 | 77.4 |  |  | 2 | 1 | 22 | 99.9 | 8.6 | 80.4 |
|  |  |  | 2 | 26 | 99.9 | 11.0 | 76.9 |  |  |  | 2 | 26 | 99.9 | 8.5 | 80.6 |
|  | EM-seq | 1 | 1 | 31 | 99.9 | 3.4 | 86.9 |  | EM-seq | 1 | 1 | 22 | 99.9 | 7.5 | 79.6 |
|  |  |  | 2 | 32 | 99.9 | 4.1 | 86.1 |  |  |  | 2 | 21 | 99.9 | 7.3 | 79.5 |
|  |  | 2 | 1 | 17 | 99.9 | 9.4 | 77.7 |  |  | 2 | 1 | 20 | 99.9 | 9.0 | 79.6 |
|  |  |  | 2 | 16 | 99.9 | 9.6 | 77.4 |  |  |  | 2 | 20 | 99.9 | 8.8 | 79.9 |
| Colon | E5hmC-seq | 1 | 1 | 29 | 99.8 | 5.0 | 85.6 | Placenta | E5hmC-seq | 1 | 1 | 28 | 100.0 | 8.8 | 79.2 |
|  |  |  | 2 | 29 | 99.8 | 4.2 | 86.3 |  |  |  | 2 | 25 | 100.0 | 8.2 | 79.8 |
|  |  | 2 | 1 | 23 | 99.8 | 9.6 | 77.7 |  |  | 2 | 1 | 21 | 100.0 | 8.0 | 80.6 |
|  |  |  | 2 | 28 | 99.8 | 11.3 | 76.7 |  |  |  | 2 | 22 | 99.9 | 9.6 | 79.0 |
|  | EM-seq | 1 | 1 | 28 | 99.8 | 4.5 | 85.9 |  | EM-seq | 1 | 1 | 22 | 99.9 | 7.3 | 80.3 |
|  |  |  | 2 | 24 | 99.8 | 4.6 | 85.8 |  |  |  | 2 | 23 | 99.9 | 7.8 | 79.8 |
|  |  | 2 | 1 | 15 | 99.8 | 10.4 | 76.7 |  |  | 2 | 1 | 25 | 99.9 | 9.9 | 78.6 |
|  |  |  | 2 | 22 | 99.8 | 10.7 | 76.8 |  |  |  | 2 | 19 | 99.9 | 8.8 | 79.3 |
| Heart | E5hmC-seq | 1 | 1 | 25 | 99.9 | 4.9 | 84.0 | Skeletal muscle | E5hmC-seq | 1 | 1 | 41 | 99.9 | 9.2 | 79.5 |
|  |  |  | 2 | 26 | 99.9 | 5.2 | 83.7 |  |  |  | 2 | 33 | 99.9 | 7.8 | 80.4 |
|  |  | 2 | 1 | 20 | 100.0 | 9.8 | 77.9 |  |  | 2 | 1 | 28 | 100.0 | 9.5 | 80.0 |
|  |  |  | 2 | 22 | 99.9 | 8.9 | 78.7 |  |  |  | 2 | 28 | 99.9 | 9.0 | 80.3 |
|  | EM-seq | 1 | 1 | 22 | 99.9 | 4.7 | 84.2 |  | EM-seq | 1 | 1 | 25 | 99.9 | 7.5 | 80.3 |
|  |  |  | 2 | 18 | 99.9 | 4.6 | 84.1 |  |  |  | 2 | 29 | 99.9 | 7.8 | 80.3 |
|  |  | 2 | 1 | 14 | 99.9 | 8.9 | 77.8 |  |  | 2 | 1 | 19 | 99.9 | 8.0 | 80.6 |
|  |  |  | 2 | 15 | 99.9 | 9.0 | 78.0 |  |  |  | 2 | 20 | 99.9 | 8.8 | 79.8 |
| Intestine | E5hmC-seq | 1 | 1 | 33 | 99.9 | 6.7 | 83.0 | Skin | E5hmC-seq | 1 | 1 | 24 | 99.9 | 7.6 | 79.4 |
|  |  |  | 2 | 31 | 99.9 | 5.8 | 83.8 |  |  |  | 2 | 26 | 99.9 | 7.9 | 79.1 |
|  |  | 2 | 1 | 24 | 99.9 | 11.1 | 77.0 |  |  | 2 | 1 | 34 | 99.9 | 12.1 | 76.5 |
|  |  |  | 2 | 23 | 99.9 | 9.3 | 78.7 |  |  |  | 2 | 22 | 99.9 | 9.4 | 78.4 |
|  | EM-seq | 1 | 1 | 20 | 99.9 | 6.0 | 83.3 |  | EM-seq | 1 | 1 | 21 | 99.9 | 7.6 | 79.1 |
|  |  |  | 2 | 18 | 99.9 | 5.8 | 82.8 |  |  |  | 2 | 25 | 99.9 | 8.3 | 78.7 |
|  |  | 2 | 1 | 21 | 99.9 | 10.7 | 77.1 |  |  | 2 | 1 | 18 | 99.9 | 10.0 | 77.3 |
|  |  |  | 2 | 17 | 99.9 | 11.3 | 76.3 |  |  |  | 2 | 16 | 99.9 | 9.2 | 77.7 |
| Kidney | E5hmC-seq | 1 | 1 | 31 | 99.9 | 5.7 | 84.0 | Spleen | E5hmC-seq | 1 | 1 | 40 | 99.9 | 7.8 | 81.1 |
|  |  |  | 2 | 23 | 99.9 | 4.7 | 84.7 |  |  |  | 2 | 45 | 99.9 | 8.1 | 80.5 |
|  |  | 2 | 1 | 30 | 99.9 | 11.5 | 76.9 |  |  | 2 | 1 | 22 | 100.0 | 10.2 | 77.9 |
|  |  |  | 2 | 19 | 100.0 | 9.2 | 78.6 |  |  |  | 2 | 32 | 99.9 | 12.4 | 76.2 |
|  | EM-seq | 1 | 1 | 38 | 99.9 | 5.6 | 83.9 |  | EM-seq | 1 | 1 | 17 | 99.9 | 6.3 | 81.3 |
|  |  |  | 2 | 25 | 99.9 | 4.8 | 84.4 |  |  |  | 2 | 20 | 99.9 | 6.4 | 81.3 |
|  |  | 2 | 1 | 18 | 99.9 | 9.3 | 78.2 |  |  | 2 | 1 | 15 | 99.9 | 10.1 | 77.1 |
|  |  |  | 2 | 20 | 99.9 | 10.4 | 77.1 |  |  |  | 2 | 20 | 99.9 | 10.5 | 77.2 |
| Liver | E5hmC-seq | 1 | 1 | 23 | 99.9 | 4.9 | 84.6 | Stomach | E5hmC-seq | 1 | 1 | 39 | 99.7 | 6.9 | 81.1 |
|  |  |  | 2 | 29 | 99.9 | 5.3 | 84.4 |  |  |  | 2 | 32 | 99.7 | 5.8 | 82.4 |
|  |  | 2 | 1 | 25 | 99.9 | 9.8 | 78.3 |  |  | 2 | 1 | 19 | 99.9 | 9.7 | 78.2 |
|  |  |  | 2 | 23 | 99.9 | 9.8 | 78.3 |  |  |  | 2 | 27 | 99.8 | 10.3 | 77.9 |
|  | EM-seq | 1 | 1 | 17 | 99.9 | 4.6 | 84.5 |  | EM-seq | 1 | 1 | 14 | 99.7 | 4.9 | 82.5 |
|  |  |  | 2 | 22 | 99.9 | 4.9 | 84.1 |  |  |  | 2 | 26 | 99.7 | 5.7 | 81.9 |
|  |  | 2 | 1 | 19 | 99.9 | 10.1 | 77.5 |  |  | 2 | 1 | 19 | 99.8 | 10.2 | 77.5 |
|  |  |  | 2 | 20 | 99.9 | 10.9 | 76.7 |  |  |  | 2 | 12 | 99.8 | 8.4 | 78.4 |
| Lung | E5hmC-seq | 1 | 1 | 34 | 99.9 | 4.6 | 85.2 | Uterus | E5hmC-seq | 1 | 1 | 29 | 99.9 | 8.3 | 80.2 |
|  |  |  | 2 | 25 | 99.9 | 4.7 | 85.0 |  |  |  | 2 | 23 | 100.0 | 8.1 | 80.1 |
|  |  | 2 | 1 | 23 | 100.0 | 10.0 | 78.2 |  |  | 2 | 1 | 28 | 99.9 | 10.3 | 78.1 |
|  |  |  | 2 | 24 | 99.9 | 9.7 | 78.4 |  |  |  | 2 | 23 | 100.0 | 9.8 | 78.3 |
|  | EM-seq | 1 | 1 | 22 | 99.9 | 5.0 | 84.7 |  | EM-seq | 1 | 1 | 21 | 99.9 | 7.7 | 80.1 |
|  |  |  | 2 | 20 | 99.9 | 4.3 | 85.1 |  |  |  | 2 | 17 | 99.9 | 7.5 | 79.8 |
|  |  | 2 | 1 | 17 | 99.9 | 9.7 | 77.8 |  |  | 2 | 1 | 23 | 99.9 | 11.3 | 76.5 |
|  |  |  | 2 | 19 | 99.9 | 10.0 | 77.6 |  |  |  | 2 | 17 | 99.9 | 9.0 | 78.2 |

**Supplemental Table S2:**
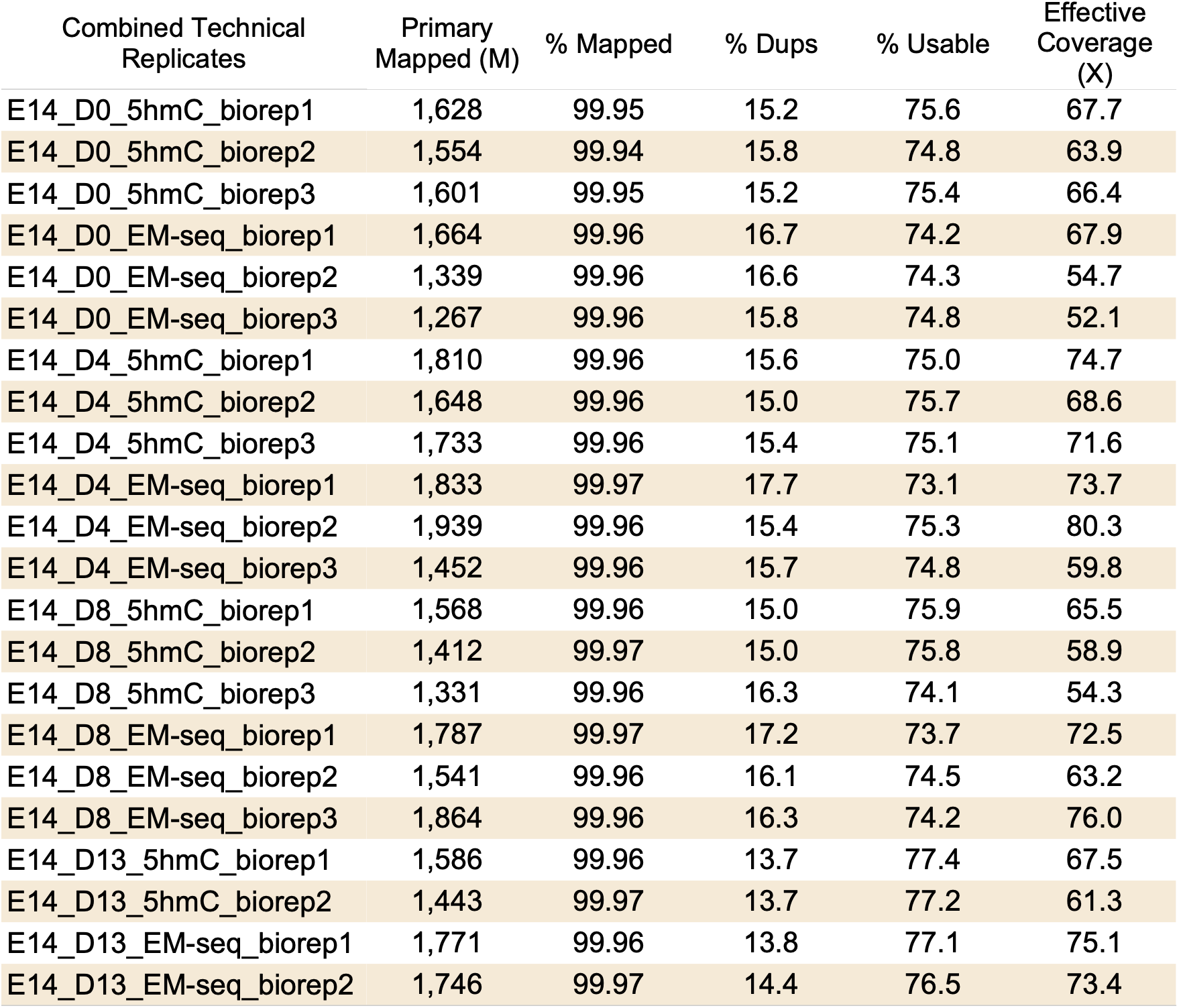
Sequencing and alignment metrics for deep-sequenced E5hmC-seq and EM-seq libraries after combining technical replicates for the neuronal differentiation time course. Theoretical coverage is calculated using the number of bases sequenced/total bases in the GRCm39 reference. (% Mapped) Reads aligned to the reference genome (GRCm39+controls); (% Dups) reads marked as duplicate by Picard MarkDuplicates; (% Usable) the set of Proper-pair, MapQ 10+, primary, nonduplicate reads used in methylation calling (SAMtools view -F 0xF00 -q 10); and (Effective Coverage) %Usable x theoretical coverage.

Supplementary Table S3: List of DETs from D0 vs D4, D0 vs D8, and D0 vs D13 and corresponding log2Foldchange.

Supplementary Table S4: List of GO:BP enrichments and their p-values corresponding to upregulated DETs from D0vsD4, D0vsD8 and D0vsD13 conditions.

Supplementary Table S5: List of GO:BP enrichments and their p-values corresponding to downregulated DETs from D0vsD4, D0vsD8 and D0vsD13 conditions.

Supplementary Table S6: List of GO:BP enrichments and their p-values corresponding to not significantly expressed DETs from D0vsD4, D0vsD8 and D0vsD13 conditions.

Supplementary Table S7: Enrichment of differential CG sites among upregulated and downregulated transcripts relative to non-differentially expressed transcripts for the same day-to-day comparison (Figure 6).

## Materials and Methods

### E14 ES cell culture and differentiation

E14 cells were grown as previously described (29) in GMEM media (Thermo Fisher Scientific) containing 10% FBS (Gemcell), 1% non-essential amino acids (NEAA, HyClone), 1% sodium pyruvate (Thermo Fisher Scientific), 50 μM beta-mercaptoethanol (Sigma-Aldrich), and 1×LIF (Millipore Sigma). To maintain the undifferentiated state, ES cells were grown on 0.1% gelatin-coated culture dishes (Stem Cell Technologies). To initiate differentiation into embryoid bodies, LIF was removed, and cells were seeded on low adherence plates (Corning) with no gelatin. The neuronal differentiation time course was done following the protocol described in (46).

### Genomic DNAs

E14 genomic DNA was extracted with the Monarch Genomic DNA Purification Kit (NEB) following the manufacturer’s instructions. Genomic DNA from fourteen C57 mouse tissues was obtained from Zyagen, USA.

### LC-MS/MS analysis

DNA samples were digested to nucleosides using Nucleoside Digestion Mix (New England Biolabs). LC-MS/MS analysis was performed by injecting digested DNAs on an Agilent 1290 Infinity II UHPLC equipped with a G7117A diode array detector and a 6495C triple quadrupole mass detector operating in the positive electrospray ionization mode (+ESI). UHPLC was carried out on a Waters XSelect HSS T3 XP column (2.1 × 100 mm, 2.5 µm) with a gradient mobile phase consisting of methanol and 10 mM aqueous ammonium acetate (pH 4.5). MS data acquisition was performed in the dynamic multiple reaction monitoring (DMRM) mode. Each nucleoside was identified in the extracted chromatogram associated with its specific MS/MS transition: dC [M+H]^+^ at m/z 228.1→112.1, d^m^C [M+H]^+^ at m/z 242.1→126.1, d^hm^C [M+H]^+^ at m/z 258.1→142.1. External calibration curves with known amounts of the nucleosides were used to calculate their ratios within the samples analyzed.

### Enzymatic 5hmC-seq and Enzymatic Methyl-seq library construction

Libraries to investigate 5hmC were generated using the E5hmC kit (NEB), whereas libraries that identified 5mC and 5hmC were generated using the EM-seq v2 kit (NEB), both following the manufacturer’s instructions. Prior to library construction, control DNAs from the kits were combined with genomic DNA following manufacturers’ recommendations. Unmethylated lambda and T4 DNA (all cytosines are 5hmC) were used for E5hmC libraries whereas unmethylated lambda and CG methylated pUC19 were used for EM-seq v2 libraries. For the E14 input titration experiments, 200 ng, 10 ng, 1 ng, 0.5 ng and 0.1 ng of DNA from undifferentiated E14 cells were used to generate E5hmC libraries. For the mouse tissue survey and E14 differentiation series, 50 ng of DNA was used to generate E5hmC and EM-seq libraries. All libraries were quantified using D1000 HS tape for TapeStation (Agilent) before Illumina sequencing (2x 150 bases, NovaSeq 6000).

### RNA-library construction

RNA-seq libraries were constructed using 100 ng of Total RNA. rRNA was depleted using the NEBNext rRNA Depletion Kit v2 (Human/Mouse/Rat) followed by library construction using the NEBNext Ultra II Direction RNA Library Prep Kit (NEB). All libraries were prepared according to the manufacturer’s recommendations. Libraries were quantified using D1000 HS tape for TapeStation (Agilent) before Illumina sequencing (2x 100 bases, NovaSeq 6000).

### Comparison of E5hmC-seq to hMeDIP

hMeDIP was performed using the Hydroxymethylated DNA Immunoprecipitation (hMeDIP) Kit (Active Motif) according to the manufacturer’s instructions with 1 µg of adult human brain DNA (Biochain) that had been fragmented to an average size of ∼350 bp using a Covaris ME220 instrument. Five replicate IP reactions were combined and purified using the Monarch® PCR and DNA Cleanup Kit (NEB) with a 5:1 binding buffer:sample ratio. After purification, duplicate libraries were prepared using 1 ng of hMeDIP DNA using the NEBNext Ultra II DNA Library Preparation Kit for Illumina. Libraries were quantified using D1000HS tape for TapeStation (Agilent) before sequencing on the Illumina platform.

Reads from hMeDIP and input control libraries were downsampled to 96M total reads, adaptor-trimmed with fastp (v0.23.4) and aligned with bowtie2 (v2.4.4) to T2T-CHM13v2.0. Duplicates were marked with picard (v3.0.0). Quality metrics were evaluated with FastQC (v0.11.9), samtools flagstats (v1.17), and picard insert sizes. Peaks were called with MACS2 against the input control.

### EM-seq and E5hmC-seq sequence data analysis

Reads were adaptor trimmed with fastp (v0.20.0) and aligned to GRCm39 (GCF_000001635.27) using bwameth (v0.2.2). Duplicates were marked with picard (v2.25.5). Quality metrics were evaluated with FastQC (v0.11.8), samtools flagstats (v1.9), picard insert sizes, and picard gc bias and checked for replicate consistency. Methyldackel (v0.6.1) was used to extract methylation information and assess methylation bias among controls. Methylation correlations were plotted with methylKit. After initial quality assessment, bams from technical replicates were combined for the remainder of the analysis, resulting in each biorep having between 1.3 and 1.9 B reads. Duplicates were re-marked using picard (v2.25.5).

Methylation information at every CG/CHG/CHH site was extracted using MethylDackel extract with the --cytosine-report option. Cytosine reports were converted to the mlml counts format, and dnmtools mlml (v1.4.2) was used to do a subtraction of 5hmC-seq methylation from EM-seq methylation (5mC + 5hmC). This gave an estimated 5mC, 5hmC, and unmethylated C proportion at each CG/CHG/CHH site.

Differentially methylated regions between days were called using defiant (v1.1.4) with all available bioreps and the following settings (coverage cutoff of 5, p-value <= 0.05, 5% difference for 5hmC and 20% difference for 5mC) to call DMRs with a minimum of 5 Cs. Counts of methylated and unmethylated C’s were estimated using the proportions from the subtracted data and the total coverage.

The mean methylation levels of called Day 0 vs Day 8 DhMRs were tracked across all days for the trajectory analysis. Hyper-DhMRs and hypo-DhMRs were clustered separately based on 5mC and 5hmC values from days 0, 4, 8, and 13 using Kmeans clustering in scikit-learn (v1.3.0).

Features were annotated using Refseq. CG islands were annotated using the UCSC annotation. Coverage across feature types was assessed using featureCounts (subread v2.0.0). The mean methylation level at features was computed using sites with at least 5 observations. Violin plots were made with seaborn (v0.12.2).

### RNA-seq analysis and data visualization

For RNA-seq analysis, adaptors were trimmed using Trim Galore, and the trimmed reads were aligned to the mm39 assembly using Salmon. We analyzed three biological replicates for each of the D0–D8 time points and two for D9 and D11, with each biological replicate comprising two technical replicates. For further analysis, we generated 3 sets of TPM matrices 1. TPM matrices for all the technical replicates; 2. TPM metrics for each bio-replicate; and 3. TPM matrices for each day point in a merge context. First and Third sets were mostly utilized for visualization of transcripts and quality control. First sets of TPM metrices with all the technical replicates were utilized to perform the PCA analysis after Variance Stabilizing Transformation (VST). plotPCA was utilized to plot technical replicates in PC1 vs PC2 plot. Similarly, a third set of TPM metrices were used for TPM based clustering. The TPM values for all the transcripts (from D0-D8, D9 and D11) were transformed to fit within a gaussian distribution like space using power transformation (yeo-johnson) using sklearn PowerTransformer package. Next, transformed data was considered for hierarchical clustering using scipy package where method was complete linkage for Euclidean distance. Finally, Heatmaps are generated using clustermap from seaborn package.

### Differential transcripts identification

Further, TPM matrices for bio-replicates were utilized for identification of Differentially expressed transcripts (DETs) using ‘local’ parameters. We calculated DETs using D0 as Control and Dn as target (n = 1,2,3,4,5,6,7,8,9,11,13) using DESeq2. Transcripts, having p-value ≤ 0.05 were considered as DE_t_, where log2FC > 1 were considered as up regulating transcripts and log2FC < 1 were considered as down regulating transcripts. Volcano plots were generated using EnhancedVolcano R library.

### Differential methylation associated with differentially expressed transcripts

To quantify relative enrichment of differentially methylated regions associated with differentially expressed transcripts, DMRs and DhMRs were intersected with transcript components: TSS region (1000 bp upstream and 100 bp downstream of TSS), first exon, exon, last exon, first intron, intron, last intron, and TES region (1000 bp downstream and 100 bp upstream of TES). Hypo and hyper DhMRs were intersected with upregulated, downregulated, and non-differential transcripts requiring at least 1bp overlap. Differential CGs among DMRs and DhMRs intersecting with a feature were counted and normalized to the number of CGs in the feature. The average number of differential CGs was obtained for each day-to-day comparison. T-tests were performed to determine if differentially methylated CGs were significantly over- or under-enriched compared to non-differentially expressed transcripts among each feature category.

## Data and Code Availability

The data underlying this article are available in the SRA submission under the BioProject PRJNA1196997. Reviewer link to the data is provided below.

https://dataview.ncbi.nlm.nih.gov/object/PRJNA1196997?reviewer=ruhkbsikjfie7bilmpa0c59ijb

Supplementary Data are available at NAR Online.

Our standard analysis pipeline for EM-seq and E5hmC-seq data is available at https://github.com/nebiolabs/EM-seq. Additional analyses specific to this paper are available at https://github.com/nebiolabs/e14_differentiation.

## Author Contributions

Daniel J. Evanich, Laura N. Blum, and Sagnik Sen contributed equally to this work. All authors reviewed and approved the final manuscript.

## Acknowledgements

We would like to thank Nan Dai for LC-MS/MS analysis as well as Rich Roberts, Tom Evans, Nicole Nichols, and Salvatore Russello at New England Biolabs Inc., for research support and encouragement.

